# COOKIE-Pro: Covalent Inhibitor Binding Kinetics Profiling on the Proteome Scale

**DOI:** 10.1101/2025.06.19.660637

**Authors:** Hanfeng Lin, Bin Yang, Lang Ding, Yen-Yu Yang, Matthew V. Holt, Sung Yun Jung, Bing Zhang, Meng C. Wang, Jin Wang

## Abstract

Covalent inhibitors are an emerging class of therapeutics, but methods to comprehensively profile their binding kinetics and selectivity across the proteome have been limited. Here we introduce COOKIE-Pro (COvalent Occupancy KInetic Enrichment via Proteomics), an unbiased method for quantifying irreversible covalent inhibitor binding kinetics on a proteome-wide scale. COOKIE-Pro uses a two-step incubation process with mass spectrometry-based proteomics to determine *k*_inact_ and *K*_I_ values for covalent inhibitors against both on-target and off-target proteins. We validated COOKIE-Pro using the BTK inhibitors spebrutinib and ibrutinib, accurately reproducing known kinetic parameters and identifying both expected and novel off-targets. The method revealed that spebrutinib has over 10-fold higher potency for TEC kinase compared to its intended target BTK. We further demonstrate that COOKIE-Pro is compatible with streamlined cysteine activity-based protein profiling (SLC-ABPP) datasets, enabling efficient conversion of competition ratios to meaningful kinetic parameters. By providing a comprehensive view of covalent inhibitor binding across the proteome, COOKIE-Pro represents a powerful new tool for optimizing the potency and selectivity of covalent drugs during preclinical development.

## Introduction

Small molecule covalent inhibitors have a rich history spanning over a century, with their mechanism initially elucidated through the study of aspirin, a classic drug. Today, this approach is emerging as a novel therapeutic modality, prized for its potency and long-lasting irreversible binding nature. A significant milestone was reached in 2013 with the FDA approval of ibrutinib, the first covalent inhibitor targeting Cys481 on Bruton’s Tyrosine Kinase (BTK), for mantle cell lymphoma (MCL) ^1^. Subsequently, covalent BTK inhibitors have evolved through several generations, each improving upon both potency and selectivity ^2^.

This progress has catalyzed the discovery of novel covalent inhibitors targeting other proteins. Notable examples include osimertinib, which targets EGFR ^3,4^, and sotorasib, which targets KRas^G12C 5,6^, a protein previously considered undruggable. Recently, advancements in chemoproteomics platform ^7–11^ together with novel covalent warhead chemistry ^12–17^ is rapidly accelerating the discovery of covalent binders and their novel targets, efficiently expanding the druggable protein universe ^18–21^. These advancements underscore the growing importance and potential of covalent inhibitors in modern drug discovery and development.

The current structural-based design strategy for covalent irreversible binders encompasses two key components: a core pharmacophore that facilitates basal reversible binding within the target protein pocket, and a covalent warhead capable of accepting nucleophilic attacks from reactive side chain groups, such as thiols on cysteine residues or phenols on tyrosine residues, to form a covalent bond. An effective covalent small molecule should possess substantial non-covalent binding affinity, ensuring sufficient residence time on the target to enhance the probability of the subsequent covalent binding event. While pursuing warheads with higher intrinsic reactivity, often referred to as ‘hot warheads’, can significantly increase the rate of covalent bond formation, this approach also elevates the risk of promiscuous off-target labeling ^7,22^. Consequently, this may lead to a higher covalent binding burden at the proteome level. Optimizing the non-covalent interactions to achieve target specificity, while maintaining a warhead with appropriate reactivity, is crucial for developing covalent inhibitors with favorable pharmacological profiles and minimal off-target effects.

Unlike reversible inhibitors whose potency can be described by dissociation constant *K*_*d*_, irreversible covalent inhibitors follow a two-step irreversible binding kinetics (Eq. 1) - the first step is the affinity-driven non-covalent binding (rate constant *k*_*on*_ and *k*_*off*_), and the second irreversible step is the covalent bond formation (rate constant *k*_*inact*_). Ranking the potency of different inhibitors is based on the second-order rate constant for covalent adduct formation, or in short, inactivation efficiency *k*_eff_, which equals the maximum rate of covalent adduct formation over inactivation constant *k*_*inact*_ /*K*_*I*_ (M^−1^ s^−1^), where *K*_*I*_ is the equilibrium constant of the first step (Eq. 2, 3). A potent covalent inhibitor must exhibit both significant intrinsic reactivity (reflected by *k*_*inact*_) and strong non-covalent binding affinity (reflected by *K*_I_). Due to the variability of the warhead intrinsic reactivity, another layer of complexity must be considered when trying to explain covalent structure-activity relationship (covSAR). Higher intrinsic warhead reactivity means a faster *K*_*inact*_, but meanwhile also increases the risk of off-target labeling, as well as a faster clearance rate *in vivo* through GSH conjugation ^23^. Optimization efforts should prioritize decreasing *K*_*l*_ to achieve tighter binding rather than switching to a more reactive warhead to push for a higher *K*_*inact*_.

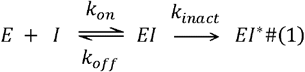

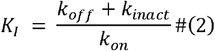

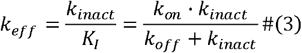

Generally, two types of methods for calculating *K*_*inact*_ and *K*_*l*_ are proposed based on whether the protein has been pre-incubated with the covalent inhibitor before adding the substrate. For the co-incubation methods, the reaction is initiated by addition of enzyme to a mixture of inhibitor and substrate, and the signal indicating product formation is recorded continuously (progression curve of time-dependent product formation for enzyme inhibition) ^24^ or obtained through a development/quenching step before readouts ^25–27^. Alternatively, pre-incubation methods take advantage of a simplified system where only the drug can interact with the protein in the pre-incubation stage, eliminating any competition from the substrate. After the pre-incubation, the substrate is added, and the fraction of remaining uninhibited proteins can be quantified by comparing the product formation rate with the uninhibited control sample ^28,29^. The choice of which experimental method to use mainly depends on the binding nature of the inhibitor (potency, or the time scale of the reaction) and the assay format (continuous reading or need post-processing). Nevertheless, all these methods require an activity-based assay for readouts of product formation.

For non-enzyme protein covalent inhibitors, *K*_*l*_ and *k*_*inact*_ can be measured using a competition-based assay with a specific chemical probe that binds to the same pocket. This method allows for the calculation of protein occupancy by the covalent inhibitor through continuous monitoring. Fluorescent polarization probes binding to KRas switch II pocket ^30^ and TR-FRET probes binding to kinase ATP pockets ^31^ have been utilized to calculate covalent inhibitor binding kinetics. Alternatively, Craven et al. developed a fluorescence-based platform using thiol-reactive probe for high-throughput kinetic analysis through quantifying the level of covalent adduct ^32,33^. However, this approach is only suitable for proteins with a single reactive cysteine, which significantly limits its potential applications. Also, one should always be cautious of any potential fluorescence artifacts, such as compound inner filtering effect especially at high concentrations, which can lead to questionable quantitative results. Recently, mass spectrometry (MS)-based quantification methods were reported either on the intact protein-adduct level ^30,34,35^ or on the peptide-adduct level ^36^. While these MS-based approaches eliminate the need for a chemical probe, prior knowledge of the off-target landscape and considerable efforts to express the target proteins are still required. As a result, measuring the kinetic parameters for all potential off-target proteins would be an extensive and labor-intensive task.

Here, we report COOKIE-Pro (<u>CO</u>valent <u>O</u>ccupancy <u>KI</u>netic <u>E</u>nrichment via <u>Pro</u>teomics), an unbiased method for irreversible covalent inhibitor binding kinetics profiling on the proteome scale, which is suitable for both enzyme and non-enzyme targets, and eliminating the need for individual protein purification by enabling direct treatment of cell lysate. As proof-of-concept, we use BTK covalent inhibitors, ibrutinib and spebrutinib, and their desthiobiotin derivatives to calculate the *k*_*inact*_ and *K*_*I*_ of the BTK inhibitors at the whole proteome scale. We also demonstrate that the data analysis method presented here is readily compatible with streamlined cysteine activity-based protein profiling (SLC-ABPP) datasets ^7^. One can easily convert the experimental condition-dependent competition ratio (CR) values of a fragment from the SLC-ABPP dataset into thermodynamic binding constants *K*_*I*_ *a*gainst the quantified cysteinome. This compatibility streamlines the data processing, enabling a more efficient translation of competition data into meaningful kinetic parameters and prioritization for downstream medicinal chemistry optimization. In addition, we propose a decision tree for the estimation of *k*_*inact*_ and *K*_*I*_ values using as few as two data points, allowing for profiling 8 compounds in each TMT-18plex run.

## Results

### Design of COOKIE-Pro Workflow

To measure the *k*_*inact*_ and *K*_*I*_ for a specific compound on the proteome scale, we need to solve two problems: 1) since MS can only identify the covalent adduct *EI*^*^ but not non-covalent intermediate *EI*, how can [*EI*^*^] be translated into *K*_*inact*_ *and K*_*I*_, and 2) how to determine [*El*^*^] for the whole proteome in a single experiment.

For the first problem, although *EI*^*^ can be monitored by MS, achieving absolute quantification requires standard peptides with known concentrations to calibrate [*EI*^*^]. This approach is not feasible on a proteome-wide scale due to the impracticality of obtaining standard peptides for every protein in the proteome. The covalent compound occupancy of a certain protein (defined as 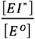) can, however, be calculated through carefully designed proteome experiments, as we will discuss below. We derived the expression for protein covalent occupancy concerning treatment time t and compound concentration [I]:

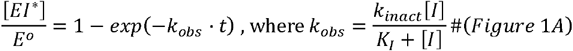

This approach allows for the determination of occupancy levels without requiring absolute quantification of the *EI*^*^ complex. Of note, this expression is based on pseudo first-order reaction kinetics without inhibitor depletion [*I*] ≫ [*E*^0^], as well as the steady state of *EI* non-covalent complex (see method mathematics part for detailed explanation).

With the expression in Figure 1A, one can obtain the *K*_*inact*_ and *K*_*I*_ values through a two-step fitting process involving multiple experimental data points with varying incubation times (t) and covalent inhibitor concentrations ([I]) (**Figure 1B**). To confirm whether the steady state approximation holds true for compounds within the realistic range of the *K*_*inact*_ and *K*_*I*_ values, we performed simulations for both low non-covalent affinity fragments (*K*_*I*_ = 10 µM) and high non-covalent affinity inhibitors (*K*_*I*_ = 10 nM) with varying *K*_*inact*_ (**Figure 1C**). The simulation results demonstrate good coherence between the theoretical and calculated *K*_*inact*_ and *K*_*I*_, indicating the approximation we used here is valid for the typical range of current covalent inhibitor kinetic parameters.

**Figure 1.**
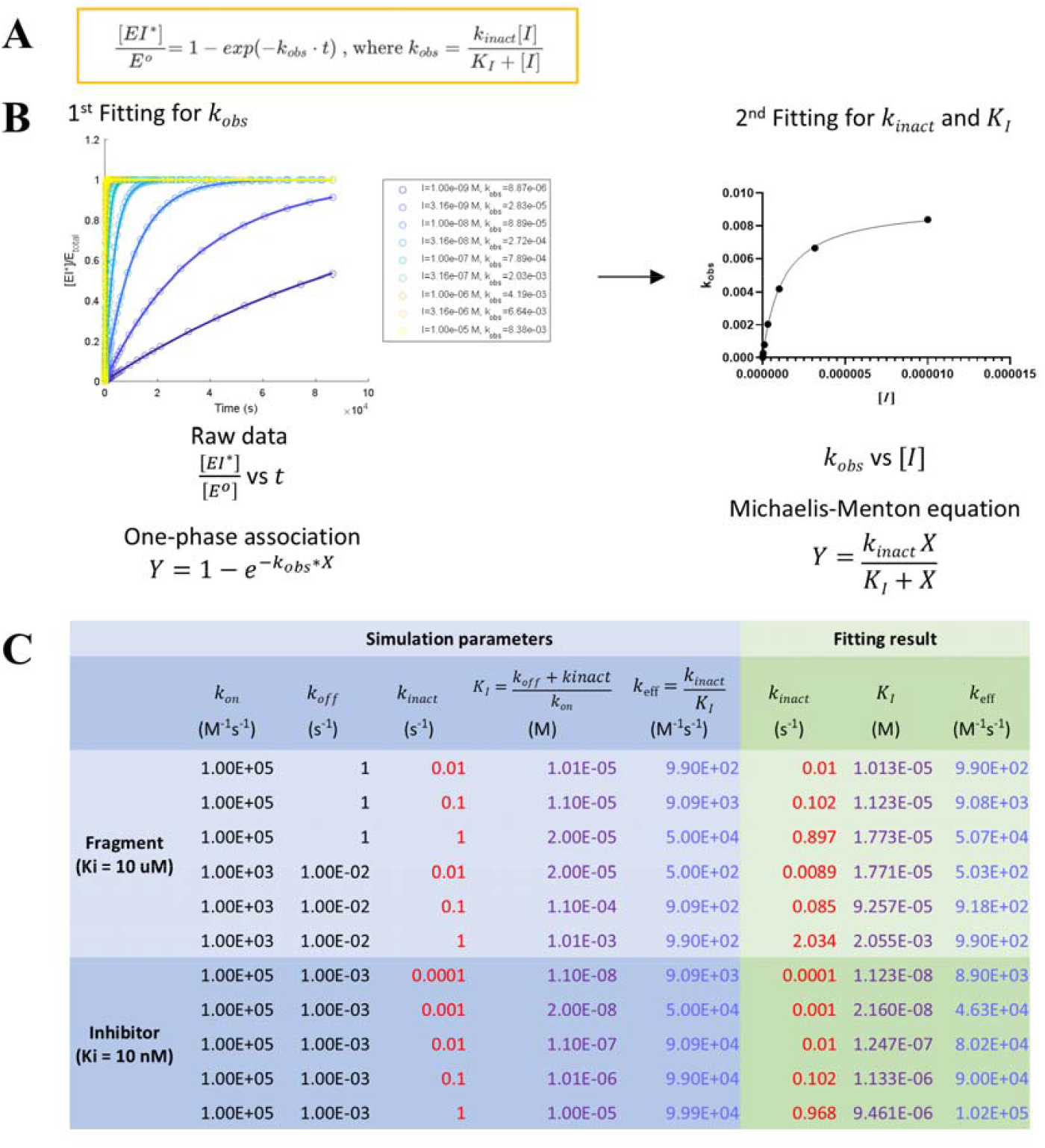
Solving *k*_*inact*_ and *K*_*I*_ using two-step model fitting and in-silico protein-ligand adduct *EI*^*^ formation kinetics simulation. **(A)** the math expression for protein covalent occupancy 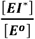 concerning incubation time ***t*** and compound concentration [***I***]. **(B)** Two-step fitting scheme to obtain ***k***_***inact***_ and KI. Protein occupancy 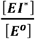 at different timepoints for a single drug concentration [***I***] will be used to obtain the apparent ***k***_***obs***_ value at this concentration through the first one-phase association fitting. Then each ([***I***]***k***_***obs***_) was plotted, and the second Michaelis-Menton fitting will be performed to obtain ***k***_***inact***_ and ***K***_*I*_ value simultaneously. **(C)** MATLAB simulation conditions and fitting results confirmed the proposed method is applicable to a wide range of compound affinities (fragments or inhibitors) and reactivity in real world settings.

With the data analysis method solved, we set out to design an experimental protocol for determining the occupancy 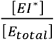 on the global proteome. The sample preparation workflow of COOKIE-Pro (Covalent <u>O</u>ccupancy <u>KI</u>netic <u>E</u>nrichment via <u>Pro</u>teomics) is shown in **Figure 2**. Generally, this is a preincubation-based method. In the first preincubation step, proteins in the cell lysate react with the compound-of-interest with varying concentration and incubation time, occupying those pockets irreversibly. However, these protein-compound adducts are not easy to quantify by untargeted MS given the substantial background proteins in the whole cell lysate. Therefore, instead of quantifying protein-compound adducts, we introduced a second incubation step using saturating concentration and incubation time of a pulldown probe followed by streptavidin bead pulldown to quantify any unreacted target proteins from the first step. This step simultaneously lyses the cell to minimize any secondary cellular response due to high concentration probe-induced oxidative stress. The desthiobiotin (DTB) probe is derived from the compound-of-interest by modifying the solvent-exposed region to mimic the covalent warhead reactivity and inhibitor affinity and selectivity. This step significantly reduces the sample complexity and improves both the depth and quantification accuracy of the protein identifications. The pulldown proteins undergo standard on-bead proteomics sample preparation workflows, including reduction, alkylation and trypsin digestion. Quantitation of the pulldown protein level among all the samples is achieved through tandem mass tag (TMT) multiplex labeling to eliminate injection-to-injection variation.

**Figure 2.**
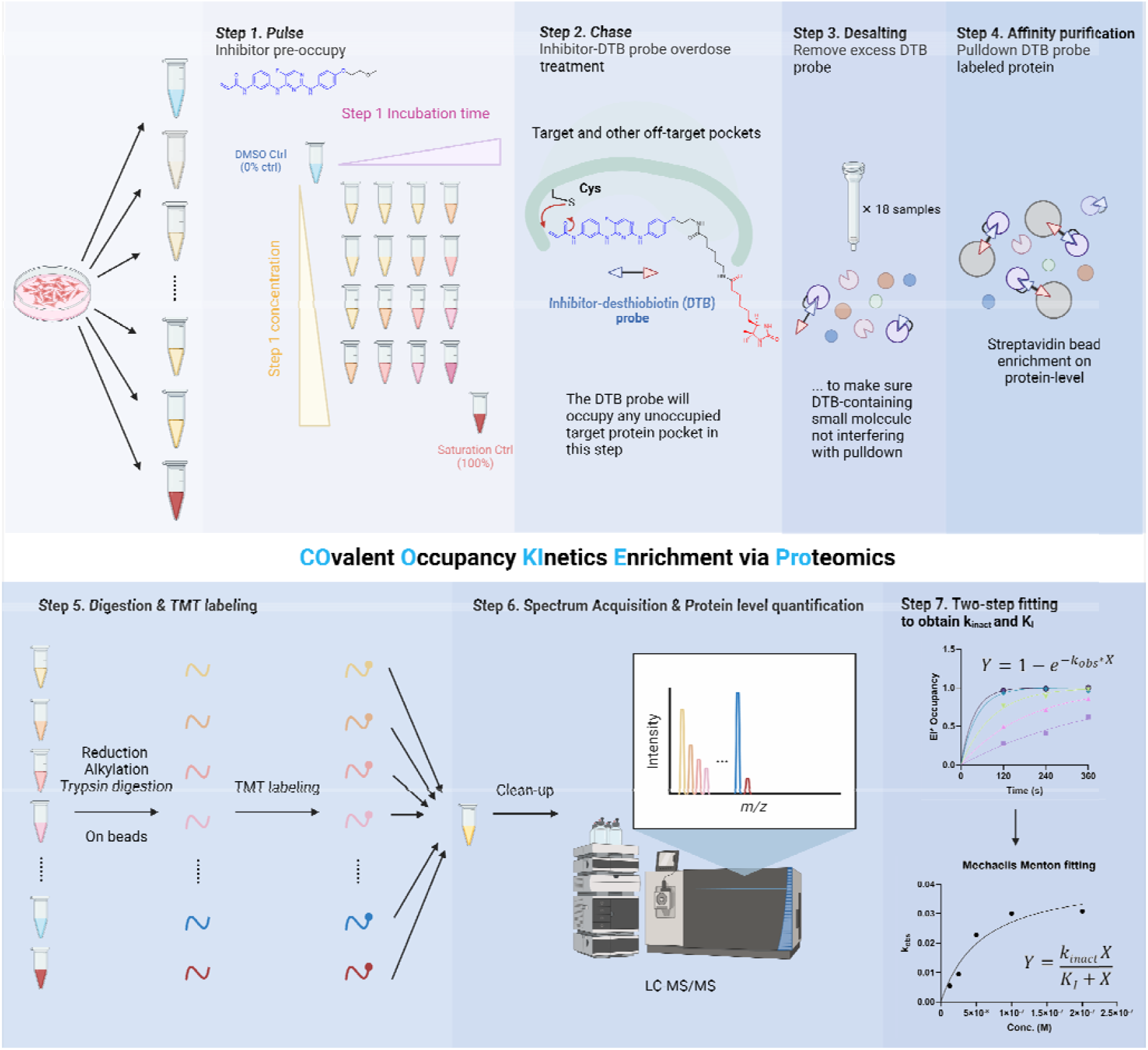
Covalent Occupancy Kinetics Enrichment via Proteomics (COOKIE-Pro) workflow using spebrutinib and its desthiobiotin probe SB-2 as an example.

Each experiment has two control samples – DMSO control and saturating control (blue and red samples in **Figure 2**). DMSO control quantifies the total probe-labeled proteome in the absence of any inhibitor competition, essentially indicates the total amount of any on-target and off-target proteins in the sample (regarded as 0% occupancy). The reported abundance ratios for a certain protein between any drug-treated sample and DMSO control sample — can be directly regarded as the unoccupied fraction of this protein (—) at this specific incubation time and concentration . It can be easily converted to occupancy for the downstrtream fitting process to obtain *k*_*inact*_ and *K*_*I*_. The saturating control, however, uses high concentration and long-enough incubation time of the covalent inhibitor to make sure that all the pockets in target proteins have been covalently pre-occupied, and the probe is no longer able to bind and pulldown these proteins. The purpose of setting up this control is to define the MS signal baseline – any ratio reported in the drug-treated groups 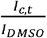 should subtract the baseline ratio 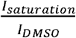, and then convert to *EL*^*^ occupancy.

### Benchmarking COOKIE-Pro with high selectivity BTK covalent inhibitor spebrutinib

To assess the entire COOKIE-Pro workflow, we chose the second generation BTK irreversible inhibitor spebrutinib (CC-292) as a benchmark compound to identify both the known target BTK and potentially undiscovered off-targets. Since substitutions on the α- and β-carbon of the acrylamide will significantly impact the warhead reactivity ^37–39^, we avoided any modification on these positions and designed the paired desthiobiotin probe SB-2 along the methoxy exit vector of spebrutinib (**Figure 3A**). Ramos cell line (B cell lymphoma) was chosen due to its high expression of the target protein BTK. To minimize any concentration interference due to cell membrane permeability but keep the proteome concentrated in the cellular environment, the cells were permeabilized with digitonin ^40^ before spebrutinib preincubation. We confirmed through microscope using trypan blue exclusion method that permeabilization happens within 30 seconds without any apparent disruption of Ramos cell morphology (data not shown). The permeabilized cells underwent two rounds of incubation: 1) For spebrutinib, we set up 5 different concentrations ranging from 25 nM to 400 nM and collected the sample from the incubation tubes every 2 minutes for 3 time points, generating a total of 15 dose-time-dependent samples plus 1 for each control sample. 2) Each sample containing ∼1.8M cells were then lysed in the NP-40 lysis buffer containing 5 µM SB-2 probe and incubated for 1 hour. The BTK amount in the pulldown samples showed good dependency on spebrutinib concentration and incubation time, while no BTK was detected in the saturation control pulldown sample (**Figure 3B**). With the sample quality confirmed, we then investigated the enrichment on the proteome level of these samples by performing TMT-based quantification.

**Figure 3.**
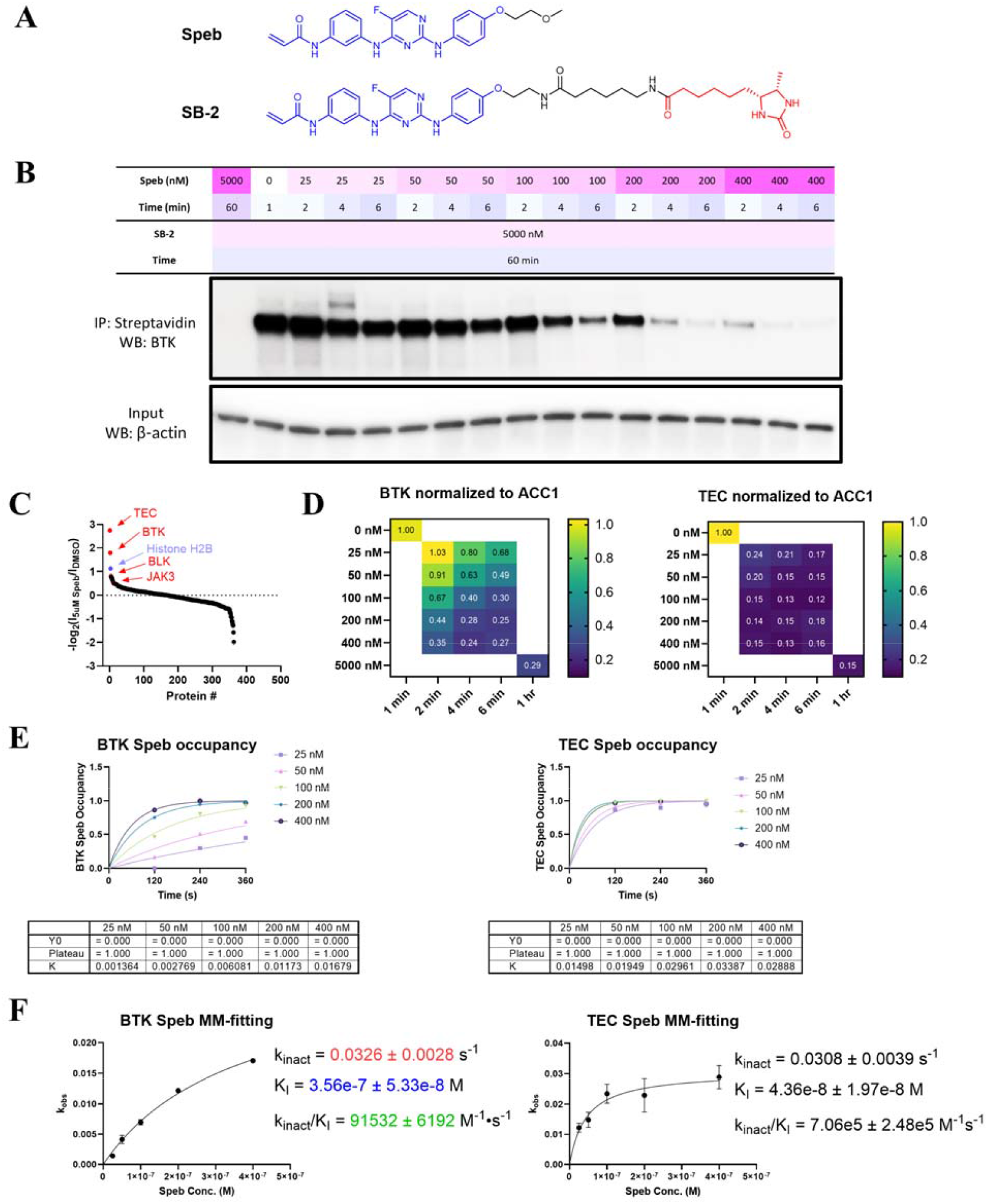
COOKIE-Pro profiling for spebrutinib in Ramos cell using SB-2 desthiobiotin probe. **(A)** Chemical structures of Spebrutinib (Speb) and the SB-2 probe. **(B)** Spebrutinib COOKIE experiment layout and the immunoblot for BTK in the pulldown samples. **(C)** Ranked proteomics -log2 transformed ACC1-normalized intensity ratio between 5 µM spebrutinib saturation sample and DMSO sample as an off-target protein list. **(D)** ACC1-normalized intensity ratio heatmaps for spebrutinib kinase off-targets across the COOKIE panel. **(E)** Representative first step progressive curve fitting for ***k***_***obs***_ at different spebrutinib concentrations against BTK and TEC. **(F)** Second step [***I***] − ***k***_***obs***_ plots fitting into Michaelis-Menton model to obtain ***k***_***inact***_ and ***K***_***I***_ values for BTK and TEC. Data are presented as mean ± SEM from three independent experiments.

Among the 412 protein groups identified, 363 protein groups are quantifiable. The top ranked protein groups not only showed high enrichment ratio between the DMSO and 5 µM saturation control groups but also across the entire concentration panel. These include TEC kinase and BTK (**Figure 3C**). Some other kinases like BLK, JAK3, NEK9 and LIMK1, only shows altered enrichment ratio between the DMSO and 5 µM saturation control groups, indicating they are insignificant off-targets of spebrutinib (**Figure S1B, Table S1**). To minimize sample-to-sample variation, we took advantage of an endogenous biotinylated protein acetyl-CoA carboxylase 1 (ACC1) which is highly and equally enriched among all the pulldown samples (**Table S1**). We normalized all the protein levels according to the ACC1 levels as an internal reference. Out of the top-ranked proteins, only BTK and TEC kinases showed dose- and time-dependent enrichment within our dosing scheme. Notably, TEC kinase demonstrated stronger binding to spebrutinib than BTK, with even the lowest concentration and shortest incubation time achieving approximately 80% occupancy (**Figure 3D**). Although only 1 peptide spectrum match (PSM) was quantified for TEC, we have verified this strong binding of spebrutinib to TEC through TEC immunoblotting of the pulldown sample (**Figure S1A**). With the occupancy data, we proceeded to COOKIE-Pro analysis using the one-phase association fitting to obtain *k*_*obs*_ (**Figure 3E**) and Michaelis-Menton fitting to obtain *k*_*inact*_ and *K*_*I*_ values (**Figure 3F**). The previously reported spebrutinib *k*_*inact*_ /*K*_*I*_ value against BTK (8.28×10^4^ M^-1^s^-1^) ^31^ is very close to our fitting result (9.15 ± 0.62 ×10^4^ M^-1^s^-1^), demonstrating the accuracy of the COOKIE-Pro workflow. We also found that spebrutinib is a very potent TEC binder which shows near 10-fold *k*_*inact*_/*K*_*I*_ against TEC (7.06 ± 2.48 ×10^5^ M^-1^s^-1^) compared with BTK. With similar *k*_*inact*_ values, the difference of inactivation potency on BTK and TEC mainly comes from the difference of the non-covalent binding affinity as indicated in *K*_*I*_.

### Optimizing COOKIE-Pro workflow with low selectivity BTK covalent inhibitor ibrutinib

With the accuracy of COOKIE-Pro confirmed, we proceeded to the next case study compound. Ibrutinib, the first generation BTK covalent inhibitor, is thought to be a more challenging case not only due to its relatively poor selectivity, but also the reported affinity value against BTK is in the tight binding range. The progression curve-based BTK kinase activity assay failed to detect any kinase activity even when ibrutinib concentration was reduced to 9 nM (**Figure S2A**). Any concentration lower than this will violate pseudo first-order reaction assumption ([*I*]should ≫ [*E*]), making it impossible to fit for kinetic parameters. The case study of ibrutinib provided valuable insights into the optimization process for COOKIE-Pro and general rules of probe design, as will be described below.

We initially designed two sets of desthiobiotin probes for ibrutinib: one has the linker attached to the β-carbon position of the acrylamide (IB-ac-DTB), and in the other set the linker was attached to the ortho-position on the piperidine ring (IB-C5-DTB and IB-C8-DTB) according to the available structure (PDB ID: 5P9I and 5P9J) ^41^. As expected, IB-ac-DTB showed weaker inhibition potency in the BTK kinase activity assay while IB-C5-DTB and IB-C8-DTB maintained a similar potency with ibrutinib (**Figure S2A**). With LC-MS, we confirmed the loss of activity of IB-ac-DTB was due to the reduced warhead reactivity (**Figure S2B**). This leads to a rule-of-thumb for pulldown probe design: avoid any modifications to the warhead moiety, even if structural information suggests that the warhead is solvent accessible. Maintaining the intrinsic reactivity of the warhead is crucial for ensuring the effectiveness of the probe.

We then switched to IB-C5-DTB and IB-C8-DTB probes. An interesting problem we encountered for IB-DTB probes was that the BTK pulldown amount no longer shows dependency on ibrutinib or spebrutinib pre-incubation, but spebrutinib-based SB-2 probe showed robust dependency on pre-incubation as expected (**Figure S3A**). We then tried lowering the concentration of the IB-DTB probes and found that only at 5 µM of either IB-DTB probe, but not at 1 µM or 0.2 µM, has significant non-specific labeling and decoupling of BTK pulldown from IB treatment was observed (**Figure S3B**). This experiment demonstrates that probes at high concentration may lose specificity and randomly react with other cysteines on the protein surface which were not pre-occupied in the first-round pre-incubation. Such non-specific labeling undermines the effectiveness of the pulldown assay and highlights the importance of optimizing probe concentration to ensure specificity.

Our probe design investigations yielded valuable insights regarding linker length. We evaluated the pulldown efficiency of IB-C5-DTB and IB-C8-DTB at varying concentrations under native pulldown conditions (**Figure S3C**). IB-C5-DTB demonstrated the ability to pulldown BTK only at 5 μM; however, the presence of β-actin in the pulldown suggested non-specific labeling. In contrast, IB-C8-DTB successfully pulled down BTK at all tested concentrations, although at 5 μM, a small fraction of BTK could not be competed off by ibrutinib preincubation, presumably due to non-specific labeling. These observations underscore the critical importance of incorporating a sufficiently long linker for effective pulldown under native conditions. Recognizing the crucial role of linker length, we hypothesized that steric hindrance between the pulldown target and streptavidin might be the primary limiting factor. To address this, we postulated that employing denaturing pulldown conditions could potentially enhance pulldown efficiency by mitigating steric hindrance and facilitating improved protein engagement. We systematically evaluated various denaturing conditions and discovered that a 0.1% SDS solution not only maintains compatibility with streptavidin beads but also significantly outperforms both native and other denaturing conditions (**Figure S4**). This finding represents a significant advancement in our probe design strategy, potentially expanding the utility and efficacy of our pulldown assays. These results highlight the importance of optimizing both linker length and pulldown conditions in the development of effective covalent probes, providing a foundation for improved target engagement and specificity in future studies.

Leveraging the insights gained during optimization, we benchmarked COOKIE-Pro for ibrutinib using the IB-C8-DTB probe (**Figure 4A**). The optimized protocol involved two sequential incubation steps with permeabilized Ramos cells: first, exposure to ibrutinib at five concentrations (12.5 nM to 200 nM) with samples collected at three time points over 6 minutes; second, cell lysis and incubation with 1 μM IB-C8-DTB probe for 1 hour. The BTK pulldown efficiency demonstrated a clear correlation with ibrutinib concentration and incubation time (**Figure 4B**). Notably, the enrichment was less pronounced compared to the spebrutinib experiment (**Figure 3B**), suggesting a higher overall covalent binding efficiency for ibrutinib. Subsequent multiplexing-based quantitative proteomics using TMTpro-18plex labeling identified 450 quantifiable protein groups. After normalization to ACC1, 346 protein groups exhibited reduced enrichment in the 5 μM ibrutinib saturation control relative to the DMSO control (**Table S2**). BTK showed the highest enrichment ratio, followed by HMGN2, BLK, H2AC20 and TEC (**Figure 4C**). Notably, BTK, BLK, and TEC are known ibrutinib off-targets in B cells ^42^, sharing cysteine homology at BTK C481. Additionally, LYN kinase was identified as a lower-ranking hit, consistent with recent SLC-ABPP findings ^10^.

**Figure 4.**
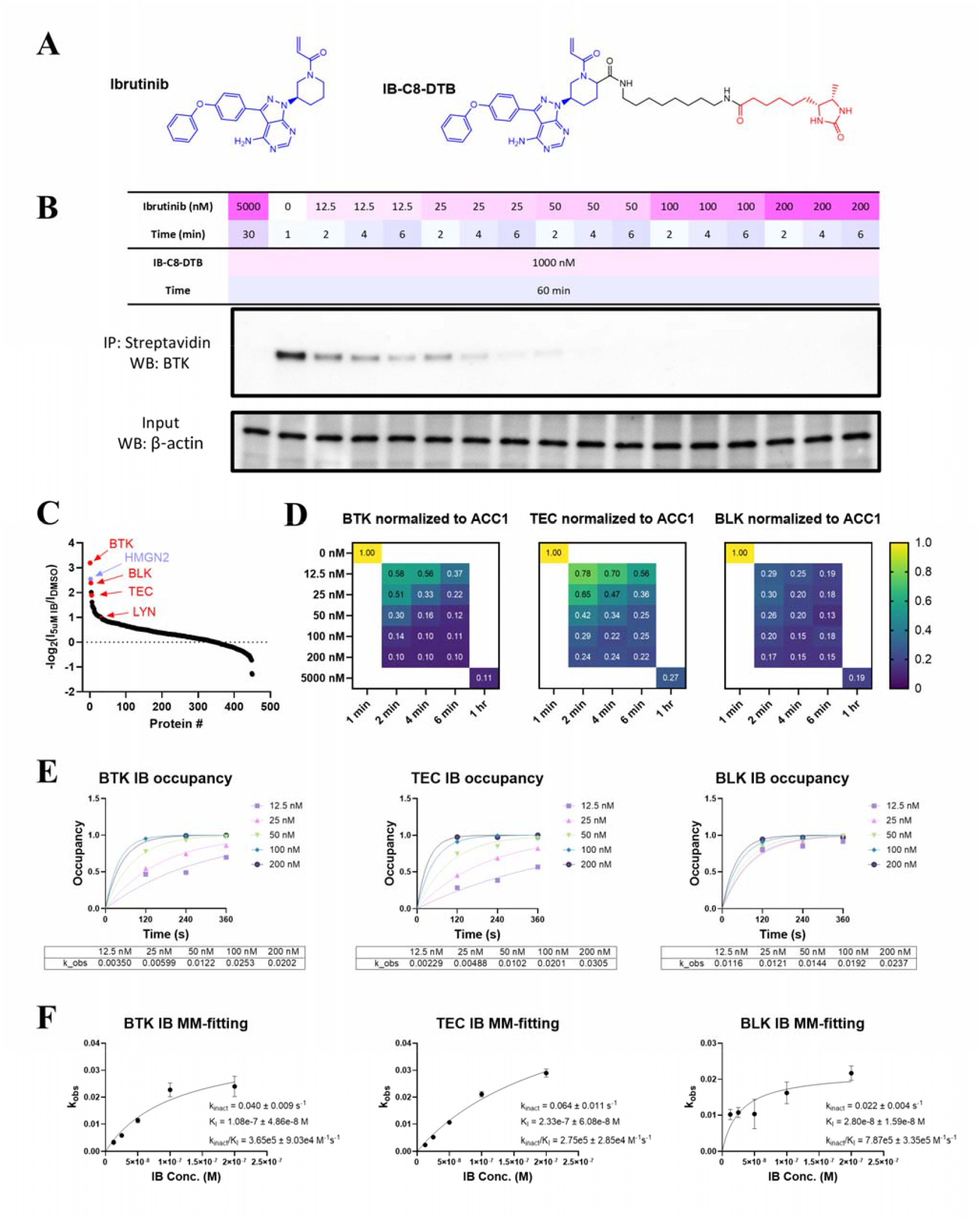
COOKIE-Pro profiling for ibrutinib in Ramos cell using IB-C8-DTB probe. **(A)** Ibrutinib (IB) and IB-C8-DTB chemical structure. **(B)** Ibrutinib COOKIE experiment layout and the immunoblot for BTK in the pulldown samples. **(C)** Ranked proteomics ACC1-normalized intensity ratio between 5 µM ibrutinib and DMSO sample as an off-target protein list. **(D)** ACC1-normalized intensity ratio heatmaps for ibrutinib kinase off-targets across the COOKIE panel. **(E)** Representative first step progressive curve fitting for ***k***_***obs***_ at different ibrutinib concentrations against BTK, TEC and BLK. **(F)** Second step [*I*] − ***k***_***obs***_ plots fitting into Michaelis-Menton model to obtain ***k***_***inact***_ (s^-1^) and (M) values for BTK, TEC and BLK. Data are presented as mean ± SEM from two independent experiments.

The three high-ranking kinases exhibited dose- and time-dependent responses to ibrutinib (**Figure 4D**). Two-step fitting of these data yielded *k*_*inact*_ and *K*_*I*_ values for each protein (**Figure 4E and 4F**). The derived parameters for BTK (*k*_*inact*_ = 0.04 s^-1^, *K*_*I*_ = 108 nM, ***k***_***inact***_ /***K***_***I***_ = 3.65×10^5^ M^-1^s^-1^) align closely with previously reported values (*k*_*inact*_ = 0.027 s^-1^, *K*_*I*_ = 54.2 nM, ***k***_***inact***_ /***K***_***I***_ = 4.77×10^5^ M^-1^s^-1^), validating the accuracy of our method. Our analysis revealed that despite similar *k*_inact_ values among the three kinases, BLK demonstrates higher susceptibility to ibrutinib (***k***_***inact***_ /***K***_***I***_ = 7.87 ×10^5^ M^-1^s^-1^) due to its stronger non-covalent affinity (*K*_*I*_ = 28 nM). Conversely, TEC exhibits weaker potency (***k***_***inact***_ /***K***_***I***_ = 2.75×10^5^ M^-1^s^-1^), attributable to its lower non-covalent affinity (*K*_*I*_ = 233 nM). These findings not only validate the COOKIE-Pro methodology but also provide valuable insights into the differential binding kinetics of ibrutinib across multiple targets, contributing to a more comprehensive understanding of its pharmacological profile.

### COOKIE-Pro strategy is compatible with SLC-ABPP dataset

High-throughput covalent fragment screening and target identification have gained considerable traction in recent years, particularly with the advancement of Affinity-Based Protein Profiling (ABPP) techniques. These methodologies enable efficient and systematic exploration of covalent interactions, facilitating the discovery of novel targets and the optimization of covalent inhibitors ^7,9,43,44^. Despite significant progress in uncovering the cysteine-ligandable proteome and identifying novel electrophilic ligands targeting these cysteines, a critical gap remains in the field: the ability to disentangle and deconvolute the contributions from non-covalent binding affinity and reactivity. This gap presents challenges in making informed decisions and managing risks associated with pursuing specific ligand-target pairs for further development.

The recently developed SLC-ABPP platform employs a thiol-reactive probe, desthiobiotin-iodoacetamide (DBIA), to enrich cysteine-containing peptides that are not covalently bonded following pre-treatment with an electrophile library ^7^. Our COOKIE-Pro workflow shares a similar underlying principle with SLC-ABPP, focusing on the enrichment and quantification of the unreacted portion of off-target candidate proteins. Given this conceptual alignment, we posit that the datasets generated from SLC-ABPP are inherently compatible with the COOKIE-Pro data analysis workflow. This compatibility offers the potential for seamless integration, enabling a more comprehensive and nuanced analysis of covalent interactions.

COOKIE-Pro relies on multiple treatment dose and time points to fit accurate kinetic parameters, but SLC-ABPP, as a screening platform, for each test compound, only single dose and single time-point data was deposited. This failed to meet the minimum requirement for obtaining an estimation of the kinetic parameters. Therefore, we assumed that fragments within the same warhead class share similar electrophilic reactivity. To estimate the reactivity parameter, we queried the CovalentInDB ^45^ for reported *k*_*inact*_ values for both nucleophilic substitution and Michael addition mechanism, and divided the data into subgroups based on warhead chemistry. The *k*_*inact*_ of the curated chloroacetamide warhead has a median of 0.0092 s^-1^, while both N-aryl and N-alkyl acrylamide have similar median *k*_*inact*_ values around 0.0025 s^-1^ (**Figure 5A-B**). This data agrees with the observation that chloroacetamide warhead fragment can label more ligandable cysteines than acrylamide fragments, providing evidence for their higher reactivity ^7,39^. We then converted the reported experimental condition-dependent competition ratios (CR) from the SLC-ABPP dataset into thermodynamic constant *K*_*I*_ values to better rank the fragments among different warhead classes. After normalization by class-specific *k*_*inact*_ values, the *K*_*I*_ values of fragments from both classes fell in the sub-mM to low mM range, with acrylamide-containing fragments showing overall tighter binding indicated by *K*_*I*_ (**Figure 5C-F, Table S3**). This is probably a survivorship bias because acrylamide-containing fragments with lower binding affinities may not show up as hits due to the lower reactivity of acrylamides compared with chloroacetamides.

**Figure 5.**
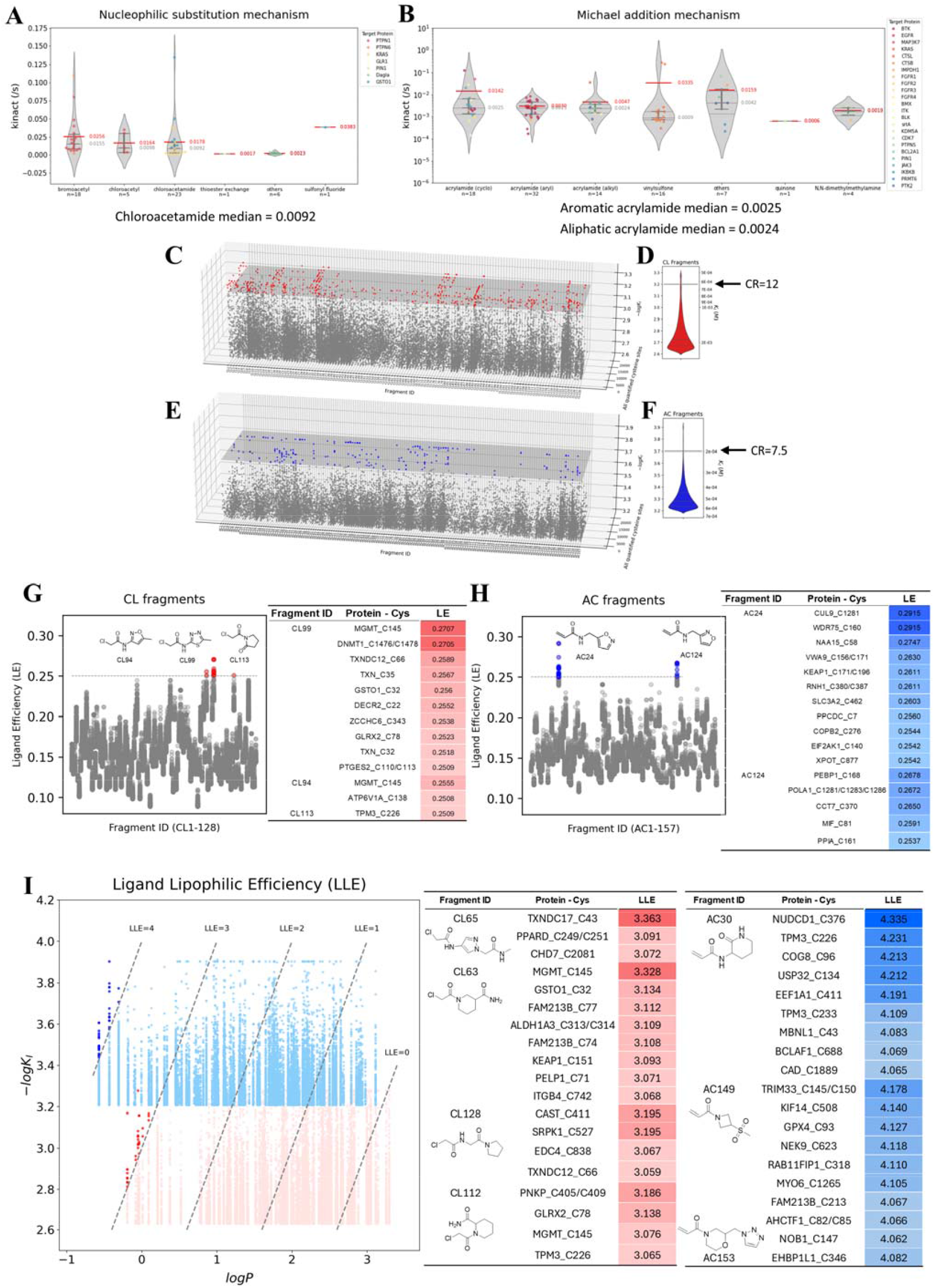
Re-analyzing SLC-ABPP data and adapting workflow to ensure COOKIE-Pro compatibility. **(A, B)** Reported values in CovalentInDB classified by reaction type and warhead subclasses. Red numbers are mean while grey numbers are median. **(C-F)** Overall ***K***_***I***_ landscapes calculated from SLC-ABPP dataset (C, E) and distribution (D, F) using COOKIE approach for chloroacetamide (C, D) and acrylamide (E, F) warheads. Arrows on panel D and F indicate the corresponding CR values at given ***K***_***I***_ values. **(G, H)** Ligand efficiency (LE) calculated from the ***K***_***I***_ values in panel C and E. **(I)** Ligand lipophilic efficiency (LLE) plot of fragment ***logP*** value versus ***logK***_***I***_ values in panel C and E. The fragment structures and targeted cysteines of highlighted dots are shown in the table.

With *K*_*I*_ values representing the non-covalent binding contribution, we calculated the ligand efficiency (LE) – a parameter that reflects the affinity contribution per atom by normalizing the affinity of compounds to their molecular size - for all the fragment-cysteine pairs (**Figure 5G-H, Table S4**). Surprisingly, with an LE cutoff of 0.25, the isoxazole moiety appeared three times among the top five fragments. Considering the two hydrogen bond acceptor atoms present in the isoxazole ring, this represents an attractive molecule pattern with high engagement efficiency that could be useful in cases where molecule weight budgets need to be considered like in the design of PROTACs or molecular glues.

Ligand-lipophilicity efficiency (LLE) is another parameter used in lead optimization that combines the potency of a compound with its lipophilicity to assess the overall quality of a drug candidate, defined as LLE = pIC_50_ – cLogP. We calculated LLE values for all the fragment-cysteine pairs (**Figure 5I, Table S5**). Fragments with higher LLE provide more room to increase lipophilicity during lead optimization without reaching an unfavorable physical profile for the drug-candidate. In contrast, low LLE indicates that the molecular activity relies excessively on hydrophobic interactions, which is generally less successful in developing effective and well-balanced drug candidates ^46^. Top ranked fragments from both classes typically have more than two hydrogen bond donor or acceptor atoms in addition to their warhead moiety, which can serve as potential enhancers for non-hydrophobic binding interactions.

Overall, we have demonstrated here the compatibility of the SLC-ABPP dataset with our COOKIE-Pro analysis workflow. This integration allows for downstream comprehensive analysis by converting experimental condition-dependent CR values into a thermodynamic inactivation constant *K*_*I*_, facilitating informed decision-making in the selection and optimization of potential drug candidates.

### Increase throughput using two data points to estimate *k*_*inact*_ and *K*_*I*_ values

In our case study, a total of 17 samples were used, including two controls and 15 dose and time titration samples. This sample size limits the throughput for screening purposes. We ask how many samples are sufficient to achieve high-throughput quantification of both *k*_*inact*_ and *K*_*I*_ without any presumption of either parameter. We were able to reduce the sample size to 7 (2 controls and 5 dose titration samples) at a single timepoint yet still obtained accurate fitting results for ibrutinib against BTK (**Figure S5**). However, sample size is still the limiting factor preventing the increase of throughput. Here, we propose a triage scheme that extracts kinetic information from a two-dose single-timepoint COOKIE fitting (**Figure 6**). This approach aims to streamline the process by providing an efficient way to obtain *k*_*inact*_ and *K*_*I*_ values with a minimal sample number, thereby enhancing throughput and making it feasible to screen a larger number of compounds or conditions.

**Figure 6.**
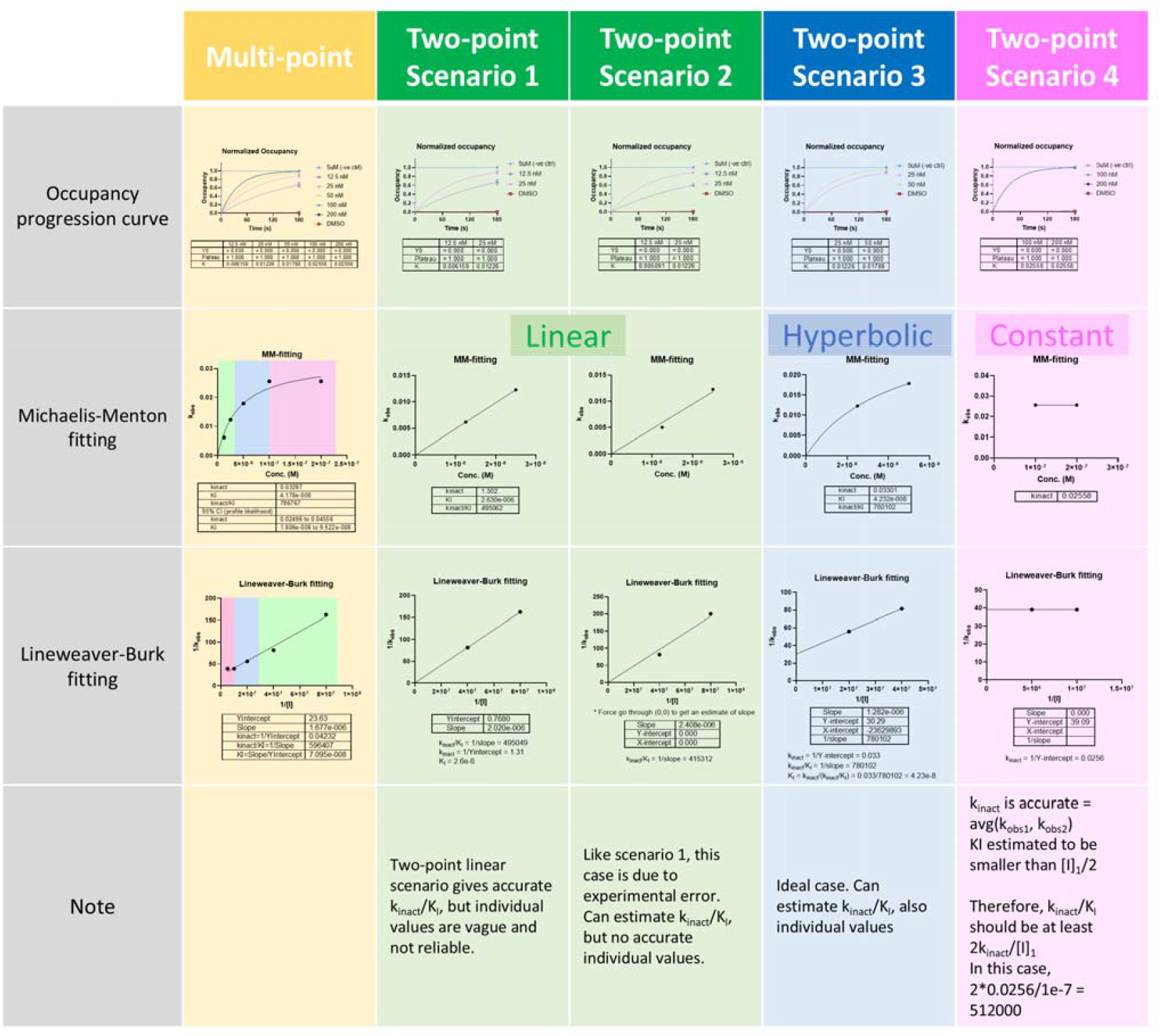
Scenarios and obtainable parameters for two-point COOKIE-Pro screening. Single timepoint with multiple concentrations are sufficient for COOKIE fitting (yellow, data from single timepoint ibrutinib COOKIE experiment **Figure S5**). Two were picked out from the five datapoints to simulate the scenarios that will be encountered in two-point COOKIE-Pro screening. **Green:** “linear phase” only gives accurate estimates of *k*_*inact*_ ***/K***_***I***_ with no individual values (fitting with linear model 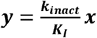). **Blue: “**curvature phase” that allows estimation of individual *k*_*inact*_ and *K*_*I*_ value (fitting with hyperbolic model 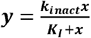). **Pink**: “plateau phase” that only ***k***_***inact***_ can be accurately estimated (fitting with constant model ***y*** = ***k***_***inact***_). The upper limit of ***K***_***I***_ is [***I***]_1_ / 2.

Since there is no pre-knowledge for the binding kinetics between the fragment to screen and its target proteins, the two concentrations would be chosen in an arbitrary manner. Therefore, the resultant datapoints on the [I]-k_obs_ plot may not always be ideal for fitting. For Michaelis-Menton fitting, the curve can be roughly divided into three stages.

1. In low concentrations where 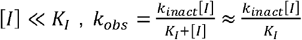, indicating a linear relationship whose slope equals to *k*_*inact*_/*K*_*I*_ (**Figure 6** green panel, scenario 1). In this scenario, minor perturbation of either data point may lead to significant changes in the individual fitting values, sometimes even failing to directly perform Michaelis-Menton fitting (as shown in **Figure 6** scenario 2). However, *k*_*inact*_ /*K*_*I*_ is rather stable and informative as it indicates the inactivation efficiency *k*_*eff*_, which can be fitted through Lineweaver-Burk plotting through origin point.
2. In middle concentrations the two points and the origin form a curvature shape (**Figure 6** blue panel, scenario 3). This is the ideal case that both individual values can be estimated through either Michaelis-Menton or Lineweaver-Burk fitting.
3. In high concentrations where 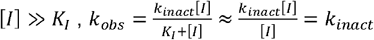. When the two datapoints on the [I]-k_obs_ plot are forming a flat line or a line with its slope < 0, it potentially means the two concentrations are already in the plateau range, therefore the average value of *k*_*obs*_ can be directly read out as *k*_*inact*_ (**Figure 6** pink panel, scenario 4). Although this scenario loses information about the concentration at which the half-maximal *k*_*obs*_ is reached (i.e., *K*_*I*_), we can still estimate that *K*_*I*_ is smaller than [*I*]/2 due to the fact that 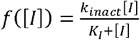 is a concave function (*f*”“c([*I*])< 0).

This triage approach using two data points offers a balance between sample economy and the accuracy of kinetic parameter estimation. It is especially advantageous in high-throughput screening settings, where the number of available samples is often a limiting factor. This method could significantly increase the throughput of screening assays, making it possible to evaluate a larger number of compounds or conditions efficiently.

### Applying two-point COOKIE-Pro strategy to profile binding kinetics for covalent fragment screening

To demonstrate the feasibility of the two-point COOKIE-Pro strategy for covalent fragment screening, 16 covalent fragments were purchased, and the screening was performed in a 96-well format using Hela cell lysate to reduce kinetic interference from membrane permeability. Cell lysates were treated with 20 or 50 μM of the fragments for 1 hour, generating 20 μg total protein per TMT channel. For each TMTpro-18plex panel, eight fragments at two concentrations were analyzed alongside two DMSO controls. This setup yielded quantification of over 10,000 desthiobiotin-iodoacetamide (DBIA)-containing peptide groups per MS injection and approximately 16,000 per panel when combining replicates (Figure S7A), generating ∼128,000 potential kinetic curves for all 8 fragments in one experiment for downstream analysis. The quantitative data was highly precise, with coefficients of variation <10% for the majority of competition ratios observed for each fragment, indicating minimal variation between replicates (Figure 7D, S7E).

**Figure 7.**
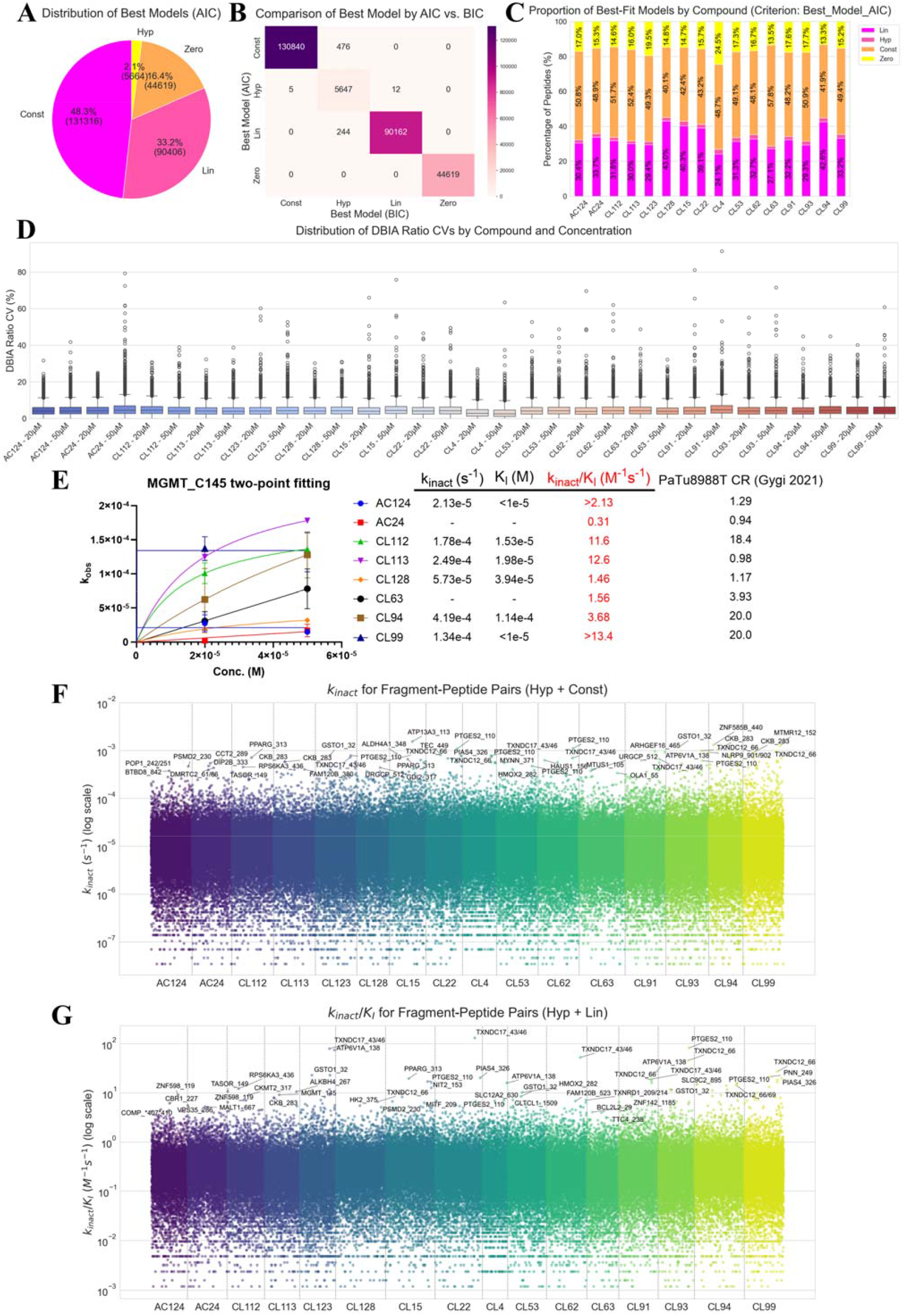
Applying two-point COOKIE-Pro strategy to profile binding kinetics for covalent fragment screening. **(A)** Pie chart showing the distribution of best-fit models (Zero, Constant, Linear, Hyperbolic) for all fragment-peptide pairs based on AIC. **<u>(B)</u>** Confusion matrix comparing the model selections made by AIC versus BIC. **<u>(C)</u>** Stacked bar chart illustrating the distribution of the four kinetic models for each of the 16 fragments screened. **(D)** Box plots showing the distribution of CVs of the DBIA ratios for each compound and concentration, indicating low variation among replicates. **(E)** A case study of two-point fitting for various fragments targeting the C145 residue of the MGMT protein. The plot shows the observed rate constant (***k***_***obs***_) versus fragment concentration, with the table summarizing the calculated kinetic parameters (***k***_***inact***_, ***K***_***I***_, ***k***_***inact***_/ ***K***_***I***_) and literature competition ratio (CR) values ^**7**^. **(F)** Scatter plot displaying the landscape of inactivation rates (***k***_***inact***_) for all peptides that fit the hyperbolic or constant models, grouped by fragment. **<u>(G)</u>** Scatter plot showing the landscape of inactivation efficiencies (***k***_***inact***_ / ***K***_***I***_) for all peptides that fit the hyperbolic or linear models, grouped by fragment.

To analyze the two-point data, we used the three models discussed in Figure 6 and selected the best fit using the Akaike information criterion (AIC) and Bayesian information criterion (BIC). AIC or BIC has been widely used in model selection decisions, with penalty for model complexity to avoid overfitting. In this study, these criteria were employed to select the best-fitting model for the two-point kinetic data. Fragment-peptide pairs with competition ratio (CR) values ≥ 1 at both concentrations were classified under a “zero model”. The AIC and BIC provided highly similar model selections (Figure 7B). Given the small sample size, we used AIC to guide model selection for downstream analysis to reduce parsimony with the hyperbolic model. Among the quantified 272,005 fragment – DBIA peptide pairs, the constant model was the most common (∼48%), followed by the linear model (∼33%). Although representing only about 2% of the total, a significant 5,664 Michaelis-Menton kinetic curves were successfully fitted using the hyperbolic model across the 16 fragments (Figure 7A). The distribution of these models varied between compounds, indicating fragment-specific kinetic behaviors (Figure 7C, S7B).

A case study on the MGMT-C145 peptide illustrates the utility of this approach (Figure 7E). Different fragments targeting this same cysteine showed distinct kinetic profiles. Fragments CL94, CL112, CL113, and CL128 fit the hyperbolic model, allowing for the determination of both *k*_*inact*_ and *K*_*I*_ . Fragments AC124 and CL99 fit the constant model, suggesting the concentrations used were in the plateau phase ([*I*] » K _*I*_). This provides an accurate estimate of kinact and a lower limit for the inactivation efficiency (*k*_*inact*_/*K* _*I*_). These kinetic parameters generally aligned with previously published CR data ^7^, with potent fragments from the original screen demonstrating high inactivation efficiencies, meanwhile provided new information about the nuance difference of intrinsic reactivity and affinity among fragments. The fitting results also showed high correlation between replicates (Figure S7C, D).

The analysis provided a broad landscape of kinetic parameters. The calculated *k*_*inact*_ values, derived from fragment – DBIA peptide pairs fitting hyperbolic and constant models, spanned several orders of magnitude from 10^-7^ to 10^-3^ s^-1^, revealing a wide range of intrinsic reactivities (Figure 7F, S7F, Table S6). Similarly, the inactivation efficiencies (*k*_*inact*_/*K*_*I*_), from pairs fitting hyperbolic and linear models, also varied widely from 10^-3^ to 10^2^ M^-1^ s^-1^, highlighting fragments with a favorable combination of non-covalent binding and chemical reactivity (Figure 7G, S7G, Table S6). The highest inactivation efficiencies (*k*_*inact*_/*K*_*I*_), on the order of 10^1^ M^−1^ s^−1^, align well with the observation that fragment-based drug discovery hits typically span a range of 10^−2^ to 10^2^ M^−1^s^−1^.^47^

Analysis of the kinetic profiling data reveals that several proteins in the thioredoxin domain-containing (TXNDC) family, such as TXNDC12 C66 and TXNDC17 C43/C46, appear as frequent, high-ranking hotspots for multiple fragments (Figure 7F,G). These proteins are involved in cellular redox regulation, a function that often relies on highly reactive, nucleophilic cysteine residues. A comparison of the kinetic parameters for these hits in Figures 7F and 7G provides insight into their binding mechanism. Cysteine sites on these TXNDC proteins rank among the most potent hits, with very high inactivation efficiencies (*k*_*inact*_/*K*_*I*_). Concurrently, these same sites also exhibit some of the highest inactivation rates (*k*_*inact*_) in the entire dataset. This strong correlation suggests that the high efficiency of these fragments against redox-prone sites is primarily driven by the high intrinsic reactivity of the cysteines (a high *k*_*inact*_), rather than by strong, specific non-covalent binding affinity (a low *K*_*I*_).

In conclusion, the application of the two-point COOKIE-Pro strategy to a covalent fragment library successfully demonstrated its utility as a high-throughput kinetic screening platform. The method efficiently generated a large and reproducible dataset, enabling the determination of kinetic parameters for thousands of fragment-cysteine interactions. By classifying interactions into distinct kinetic profiles, this approach provides a more nuanced understanding than traditional single-point screening in SLC-ABPP. The ability to decouple reactivity from affinity at scale offers a significant advantage for medicinal chemistry, allowing for a more informed prioritization of hits. This streamlined workflow enhances the efficiency of covalent drug discovery by providing the detailed mechanistic insights necessary to guide the rapid optimization of fragments into selective and potent drug candidates.

## Discussion

The evolution of covalent inhibitor development underscores the significance of this novel drug class in targeting difficult-to-drug proteins, with notable advancements in designing inhibitors that balance potency, selectivity, and safety ^34–36,45^. The kinetics parameters of covalent inhibitors, *k*_*inact*_, *K*_*I*_, and *k*_*eff*_, provide valuable insights into balancing potency with selectivity but are currently limited by the prior knowledge of on-target and off-target proteins and low-throughput assays for determining these parameters against these proteins.

The COOKIE-Pro method, introduced in this study, represents a significant advancement in profiling covalent inhibitor binding kinetics unbiasedly on a proteome-wide scale. This method addresses two major challenges in covalent inhibitor research: the quantification of covalent adducts and the determination of binding kinetics across the proteome. COOKIE-Pro utilizes a preincubation-based approach combined with MS-based proteomics, enabling the calculation of *k*_*inact*_ and *K*_*I*_ for covalent inhibitors across a broad range of proteins. The use of a desthiobiotin probe in the second incubation step and subsequent pulldown significantly reduces sample complexity, thereby enhancing the sensitivity of protein detection and addressing challenges posed by low-abundance targets. Although the DBIA probe used in SLC-ABPP can nonspecifically react with solvent-exposed cysteinome, it is not suitable for the COOKIE-Pro protein-level enrichment workflow – proteins with multiple DBIA reactive cysteines will still be enriched even if one inhibitor-specific cysteine is pre-occupied. While the DBIA workflow could function if the proteome is digested into peptides prior to streptavidin enrichment, it may still face challenges in identifying overly short or long tryptic peptides, as seen with the BTK C481-containing peptide (21 residues). Our protein-level enrichment approach better meets the need for deep profiling of potent inhibitor binding targets.

The validation of COOKIE-Pro with high selectivity BTK inhibitor spebrutinib demonstrated the method accuracy and reliability. The subsequent application to ibrutinib provided further insights into the method utility. Despite the lower selectivity of ibrutinib compared to spebrutinib, COOKIE-Pro successfully identified and quantified known off-targets in the Ramos cells, such as BLK and TEC ^42^, providing valuable kinetic parameters (*k*_*inact*_ /*K*_*I*_) that are consistent with previous reports for BTK protein ^31,48^. This consistency underscores the robustness of COOKIE-Pro in accurately profiling covalent inhibitor interactions, even for compounds with complex binding profiles.

The ibrutinib case study highlighted several important considerations for probe design and experimental conditions, including the impact of warhead reactivity, choosing a proper concentration to minimize non-specific labeling, and the effect of linker length on pulldown efficiency. The optimization process revealed that maintaining the intrinsic reactivity of the warhead is crucial for effective probe design, and that switching from native to denaturing pulldown workflow can minimize steric hindrance and improve pulldown efficiency. The concentration of probe used should be high enough to out-compete all the remaining non-covalent binding with the inhibitor as quickly as possible. However, using too high a probe concentration should be avoided, as non-specific labeling could occur. Large step dilution from the first to the second incubation is theoretically recommended over using saturating probe concentration, but in practice, it may reduce the identified protein numbers due to less input material amount.

The combination of SLC-ABPP with the COOKIE approach presents an opportunity to enhance the efficiency and throughput of kinetic profiling in drug discovery and chemical biology. While SLC-ABPP typically provides an experiment-dependent competition ratio (CR) as a measure of inhibitor potency, integrating the COOKIE approach aims to quantify intrinsic kinetic parameters *k*_*inact*_ and *K*_*I*_, which are fundamental characteristics of the enzyme-inhibitor interaction and independent of the specific experimental setup. Due to limitations in the published dataset containing only one datapoint, a surrogate method is used to assume an identical *k*_*inact*_ for fragments within the same warhead class and calculate the *K*_*I*_ values for each fragment-cysteine pair.

This study further demonstrated the feasibility of a high-throughput screening application by adapting the workflow into a two-point COOKIE-Pro strategy to profile a covalent fragment library. This approach moves beyond the single, experiment-dependent competition ratio (CR) offered by platforms like SLC-ABPP by quantifying the intrinsic kinetic parameters *k*_*inact*_ and *K*_*I*_ during fragment-based drug screening. The screening of 16 fragments generated a rich dataset of over 100,000 potential kinetic curves. By fitting data to distinct kinetic models, this approach provides a more nuanced understanding than single-point screening. A key insight from this analysis was the ability to decouple chemical reactivity from non-covalent affinity. For instance, frequent hits on thioredoxin domain-containing (TXNDC) proteins were found to have high inactivation efficiencies (*k*_*inact*_ /*K*_*I*_). Our analysis revealed this was primarily driven by the high intrinsic reactivity of these redox-prone cysteines (a high *k*_*inact*_) rather than specific binding affinity. Furthermore, the observed inactivation efficiencies, with top hits on the order of 10^1^ M^−1^s^−1^, are consistent with the potency range typically expected for fragment-based hits. This ability to generate detailed mechanistic information at scale is a critical advantage for guiding medicinal chemistry efforts.

It is important to note that while the two-point COOKIE method offers significant advantages in terms of sample efficiency and throughput, there are important statistical limitations and considerations. With only two data points used to estimate both *k*_*inact*_ and *K*_*I*_, the fitting process is prone to overfitting, potentially resulting in parameter estimates that are overly sensitive to small variations in the data. This can lead to inaccurate or non-reproducible results. The limited data points also make it difficult to accurately estimate the uncertainty or confidence intervals around the fitted parameters. In real screening situations, it is important to use *k*_*obs*_ values fitted from multiple biological replicates to minimize the *k*_*inact*_/*K*_*I*_ fitting error. Misinterpreting the scenario (whether the data points fall into the low, middle, or high concentration range) can lead to incorrect conclusions about the kinetic parameters, particularly if the chosen concentrations are close to *K*_*I*_. Also, when selecting the two concentrations, a low-dose and high-dose combination increases the chance of fitting into Scenario 3, from which more information could be inferred.

In summary, the introduction of COOKIE-Pro represents a significant advancement in covalent inhibitor research, offering a powerful tool for profiling inhibitor binding kinetics across the proteome. The method’s successful application to both highly selective and less selective inhibitors demonstrates its versatility and potential for broad application in drug discovery and development. The successful extension of this strategy to a high-throughput, two-point format for fragment screening further underscores its value in early-stage drug discovery. The ability to quantitatively assess these interactions on a proteome-wide scale opens new avenues for understanding the full spectrum of covalent inhibitor activity and optimizing drug design for better therapeutic outcomes. Future studies could explore the application of COOKIE-Pro to other classes of covalent inhibitors and in different cell lines and tissues, integrating the binding kinetics parameters across the whole proteome into PK/PD modeling and toxicology predictions. We also envision with the development of data-independent acquisition (DIA) high-throughput proteomics, the theoretic framework of COOKIE-Pro can be readily applied to covalent fragment screen to prioritize the target and fragment pairs.

## Methods

### Mathematics

Consider a 2-step irreversible inhibitor *I* binding to its target protein *E*, the reaction can be described as:

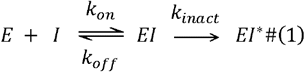

The equilibrium constant *K*_*I*_ (eq. 2) for the first step was defined based on the steady state approximation of *EI* (eq. 4):

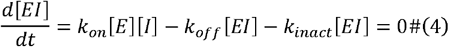

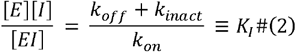

The rate constant for *EI*^*^ adduct formation:

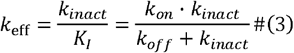

Mass law for total target protein *E*^0^ and non-covalent protein ε:

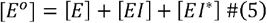

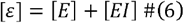

Assuming the reaction follows pseudo first-order reaction kinetics without inhibitor depletion, the unbound inhibitor concentration [*I*] is a constant value unaffected by enzyme binding.

Since proteomics experiment can only quantify [*EI*^*^], the goal is to define 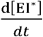.

The formation rate of [*EI*^*^] equals the consumption rate of [*ε*]:

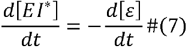

On the other hand, by rearranging Eq. 2 with 6, we solve for [*EI*]:

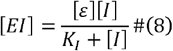

Therefore, the formation rate of [*EI*^*^] can be expressed as:

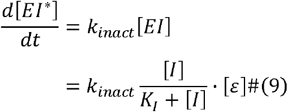

Combining Eq. 7 and 9, we get a differential equation for [*ε*]:

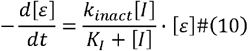

This equation can be solved by rearranging and integrating on both sides:

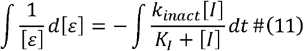

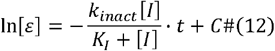

Here, we took advantage of the integration constant *c*, let *c*= ln[*E*^°^]. After rearranging, we got:

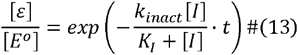

Substitute [*ε*] with [*E*^°^] − [*EI*^∗^] and rearrange, we finally got the equation in Figure 1A.

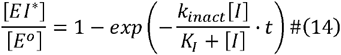

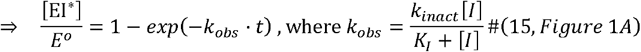

### MATLAB simulation

MATLAB (version R2022b) and the SimBiology package (v6.4) were used to build and simulate the covalent inhibition model as shown in Eq. 1. The system was built as described in the following 4 ordinary differential equations (ODEs):

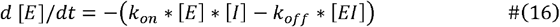

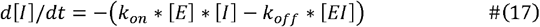

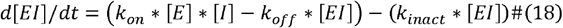

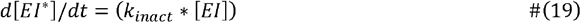

The initial value of [*E*] was set at 1e-10 M, [*I*] titrating from 1e-9 to 1e-5 M with 9 steps on the log scale. Both [*EI*] and [*EI*^∗^] were initially set as 0. The values of *k*_*0n*_, *k*_*0ff*_, and *k*_*inact*_ were set as each condition in Figure 1C. The simulation stop time was set at 86,400 seconds. ODE15s solver was used for all the simulation.

The simulated datapoints of [*EI*^∗^] were exported and used to fit for *k*_*0bs*_ values using *lsqcurvefit* function in the MATLAB. The fitting model = @(params, x) params(1) + (params(2) -params(1)) * (1 - exp(-params(3) * x)), where params(1)=0, params(2)=1e-10, and params(3) is *k*_*obs*_ to be fitted. The fitted *k*_*obs*_ values and their corresponding [*I*] were used for the second round Michaelis-Menton fitting in GraphPad 10.2 to generate the individual *k*_*inact*_ and *K*_*I*_values.

#### Chemistry

#### Synthesis of SB-2

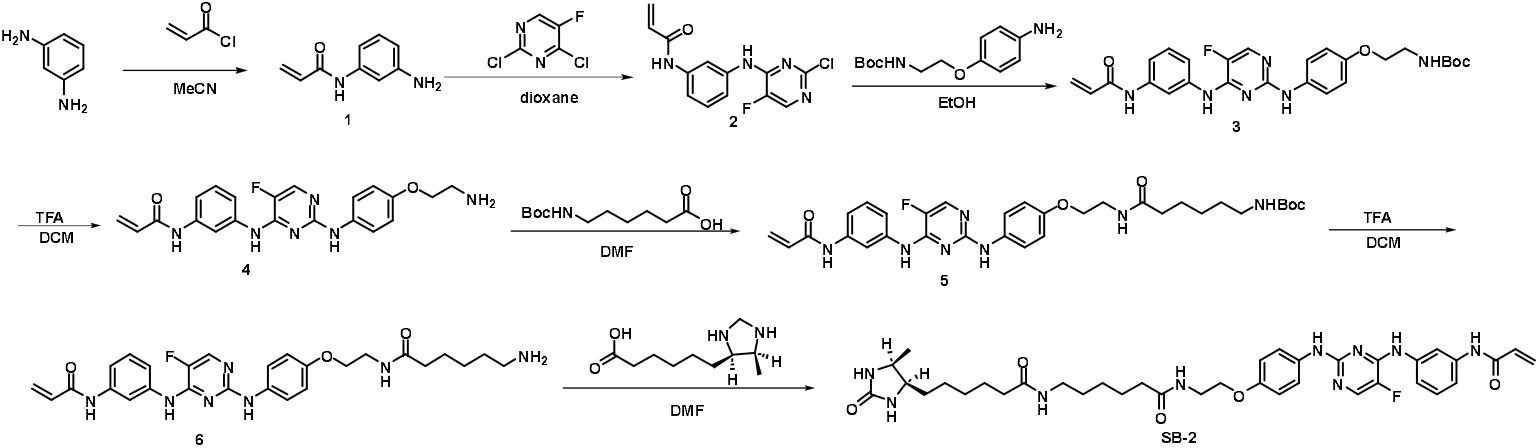

A mixture of *m*-phenylenediamine (487 mg, 4.51 mmol), acetonitrile (9 mL) and sodium bicarbonate (1.01 g, 12.00 mmol) were cooled to 0°C in an ice bath, then acryloyl chloride (272 mg, 3.00 mmol) was added dropwise over 10 min, and the resulting mixture was allowed to warm to room temperature and stirred for another 30 min. The reaction mixture was then poured into water (60 mL), filtered and washed with water to give the crude intermediate **1**.

To a solution of 2,4-dichloropyrimidines (410 mg, 2.45 mmol) in 1,4-dioxane was added intermediate **1** (332 mg, 2.04 mmol) and DIPEA (396 mg, 3.07 mmol). The reaction mixture was stirred at 80 °C for 4 h. After being cooled to room temperature, the reaction mixture was added to water (30 mL). The precipitate was filtered, and the filtered cake was rinsed with additional cool water and dried in a vacuum oven to give the intermediate **2**, which were used without further purification.

To a solution of **2** (200 mg, 0.683 mmol) and *tert*-Butyl 2-(4-aminophenoxy)ethylcarbamate (276 mg, 1.09 mmol) in ethanol (4 mL) was added glacial acetic acid (20.5 mg, 0.342 mmol), and the reaction mixture was stirred in a sealed tube for 16 h at 90 °C. The reaction mixture was cooled, concentrated under reduced pressure. The residue was quenched with a 10% sodium bicarbonate solution (4 mL) and extracted with ethyl acetate (3x10 mL). The combined ethyl acetate was washed with water (10 mL), brine (10 mL), dried over Na_2_SO_4_ and concentrated under reduced pressure. The crude residue was further purified by column chromatography to get intermediate **3** (165 mg, 47%).

To a solution of **3** (165 mg, 0.324 mmol) in DCM (1 mL) was added TFA (0.5 mL). The reaction mixture was stirred for 2 h. The mixture was concentrated under reduced pressure to get intermediate **4** (133 mg, 73 %).

To a solution of **4** (38.4 mg, 0.094 mmol) and Boc-6-aminohexanoic acid (26.1 mg, 0.113 mmol) in DMF (4 mL) was added HATU (71.5 mg, 0.188 mmol) and DIPEA (48.6 mg, 0.376 mmol). The reaction mixture was stirred at room temperature overnight. The mixture was concentrated under reduced pressure and was further purified by column chromatography to get intermediate **5** (37 mg, 63%).

To a solution of **5** (37mg, 0.324 mmol) in DCM (1 mL) was added TFA (0.5 mL). The reaction mixture was stirred for 2 h. The mixture was concentrated under reduced pressure to get intermediate **6** (26.9 mg, 97 %).

To a solution of **6** (26.9 mg, 0.052 mmol) and *d*-desthiobiotin (12.1 mg, 0.0.057 mmol) in DMF (1 mL) was added HATU (39.2 mg, 0.103 mmol) and DIPEA (33.3 mg, 0.258 mmol). The reaction mixture was stirred at room temperature overnight. The mixture was concentrated under reduced pressure and was further purified by column chromatography to get **SB-2** (31 mg, 84%).

#### Synthesis of IB-ac-DTB

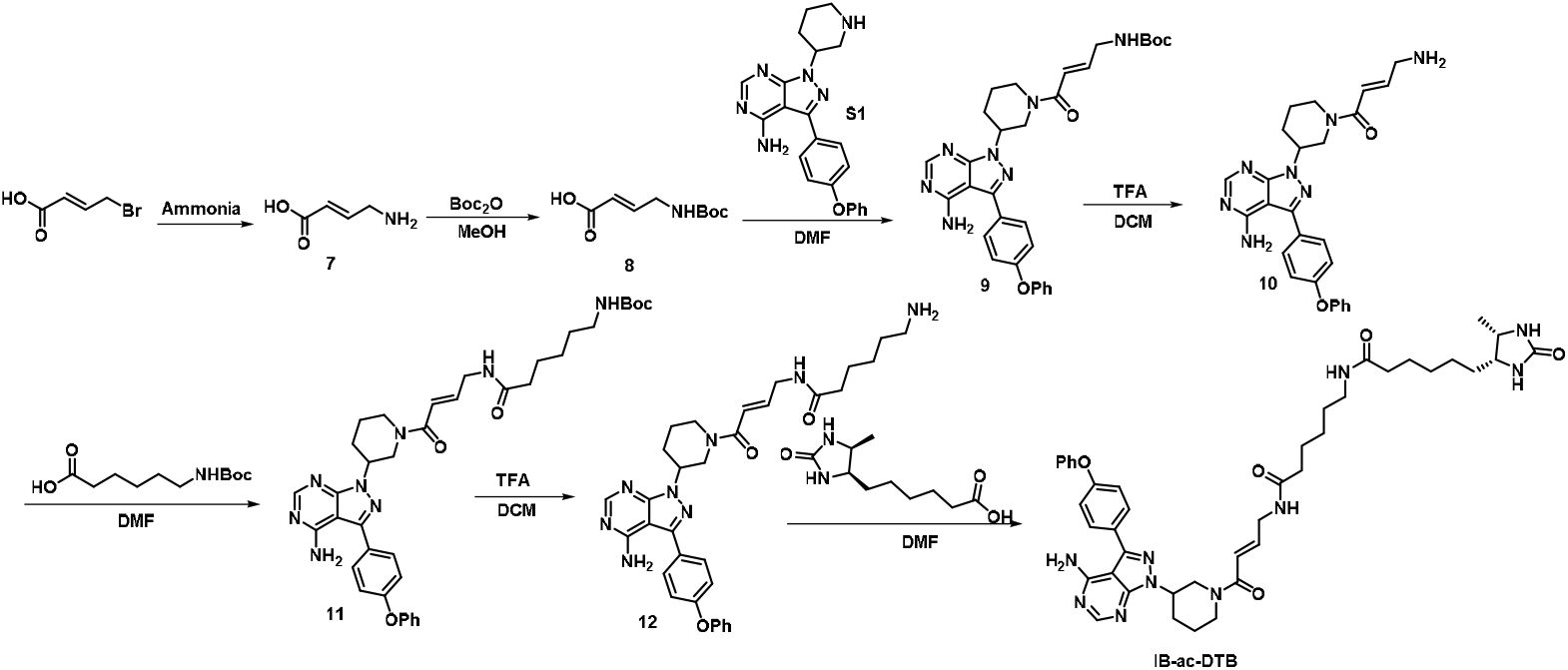

To a vial were added (E)-4-bromobut-2-enoic acid (100 mg, 0.6 mmol) and aqueous ammonia (1 mL) and was stirred at room temperature overnight. The mixture was concentrated under reduced pressure to dryness to yield the intermediate **7** (57 mg, 93%), which was used in the next step without purification.

To a vial containing a solution of **7** (57 mg, 0.56 mmol), Na_2_CO_3_ (120.9 mg, 1.12 mmol), THF (2 mL), and H_2_O (2mL) was added Boc_2_O (241.8 mg, 1.12 mmol). The reaction mixture was stirred at room temperature overnight. Then the mixture was extracted with EtOAc and the pH of the aqueous layer was adjusted to 2 with HCl (1M). The aqueous layer was extracted with EtOAc, washed with brine (3X), dried over anhydrous Na_2_SO_4_, filtered, and concentrated under reduced pressure to yield the intermediate **8** (96.9 mg, 86%).

To a solution of **S1** (185.2 mg, 0.48 mmol) in 10 mL DMF was added HOBT (100.7 mg,0.72 mmol), EDC (138.0 mg, 0.72 mmol), DIPEA (124.1 mg, 0.96 mmol) and **8** (96.9 mg, 0.48 mmol), successively. The reaction was stirred at room temperature overnight prior to addition of 8 mL water and then extracted with ethyl acetate (2 × 50 mL), the combined organic phase was washed with brine, dried over anhydrous Na_2_SO_4_ and concentrated under reduced pressure. The residue was purified by flash column to give intermediate **9** (217.7 mg, 76%).

To a solution of **9** (217.7 mg, 0.365 mmol) in DCM (5 mL) was added TFA (2 mL). The reaction mixture was stirred for 1h. The mixture was concentrated under reduced pressure to give intermediate **10** (171.4 mg, 100%), which was used in the next step without purification.

To a solution of **10** (171.4 mg, 0.365 mmol) and Boc-6-AMinocaproic acid (101.3 mg, 0.438 mmol) in DMF (5 mL) was added HATU (277.6 mg, 0.73 mmol) and DIPEA (188.7 mg, 1.46 mmol). The reaction mixture was stirred at room temperature overnight. The mixture was concentrated under reduced pressure and was further purified by column chromatography to get intermediate **11** (107.2 mg, 42%).

To a solution of **11** (107.2 mg, 0.157 mmol) in DCM (2 mL) was added TFA (1 mL). The reaction mixture was stirred for 1 h. The mixture was concentrated under reduced pressure to give intermediate **12** (91.5 mg, 100%), which was used in the next step without purification.

To a solution of **12** (91.5 mg, 0.157 mmol) and *d*-desthiobitin (40.4 mg, 0.188 mmol) in DMF (2 mL) was added HATU (100 mg, 0.314 mmol) and DIPEA (81.2 mg, 0.628 mmol). The reaction mixture was stirred at room temperature overnight. The mixture was concentrated under reduced pressure and was further purified by column chromatography to give **IB-ac-DTB** (107.2 mg, 42%).

#### Synthesis of IB-C5-DTB and IB-C8-DTB

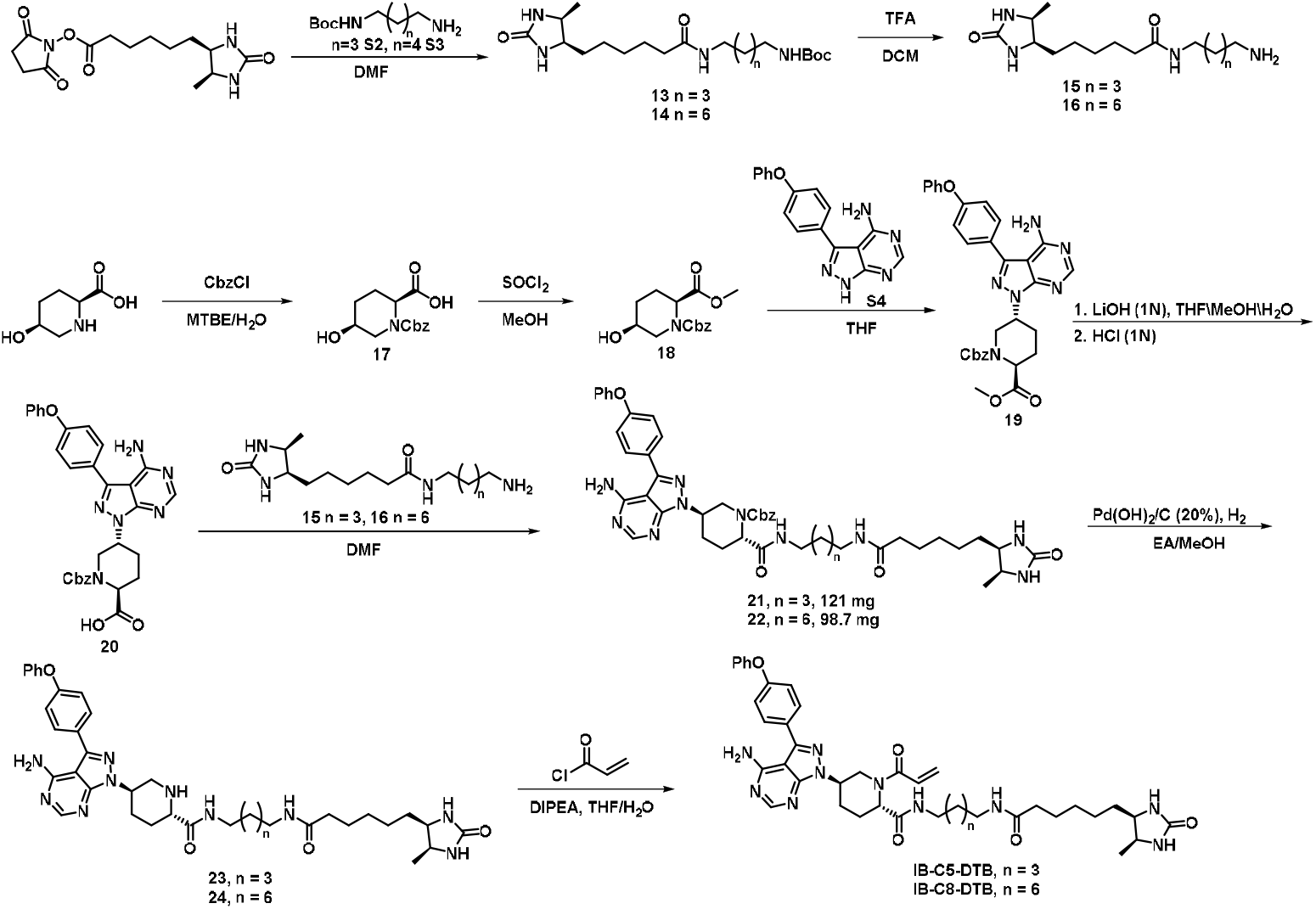

To a solution of **S2** (80.9 mg, 0.4 mmol) in DMF (2 mL) was added *d*-dethiobiotin-NHS (62.3 mg, 0.2 mmol) and DIPEA (77.5 mg, 0.6 mmol). After the solution was stirred overnight at room temperature, the solvent was removed by rotary evaporation. The residue was further purified by column chromatography to give intermediate **13** (94.7 mg, 95%).

To a solution of **S3** (97.7 mg, 0.4 mmol) in DMF (2 mL) was added *d*-dethiobiotin-NHS (62.3 mg, 0.2 mmol) and DIPEA (77.5 mg, 0.6 mmol). After the solution was stirred overnight at room temperature, the solvent was removed by rotary evaporation. The residue was further purified by column chromatography to give intermediate **14** (97.9 mg, 91%).

To a solution of **13** (94.7 mg, 0.19 mmol) in DCM (2 mL) was added TFA (1 mL). The reaction mixture was stirred for 1h. The mixture was concentrated under reduced pressure to give intermediate **15** (75.7 mg, 100%), which was used in the next step without purification.

To a solution of **14** (97.9 mg, 0.18 mmol) in DCM (2 mL) was added TFA (1 mL). The reaction mixture was stirred for 1h. The mixture was concentrated under reduced pressure to give intermediate **15** (79.3 mg, 100%), which was used in the next step without purification.

To a solution of *cis*-5-hydroxy-L-pipecolic acid (145.2 mg, 1 mmol) and potassium carbonate (172.8 mg, 1.25 mmol) in MTBE (1 mL) and H_2_O (5 mL) was added benzyl chloroformate (187.6 mg, 1.1 mmol) dropwise at 15 °C. After stirring for 3 h, the mixture was abstracted with MTBE (2 mL). Then the organic phase was discarded, and the aqueous phase was acidified with concentrated HCl until pH=2 at 0 °C. The acidified aqueous phase was extracted with ethyl acetate (3 × 10 mL), and the combined organic phase was washed with brine, dried over anhydrous Na_2_SO_4_ and concentrated under reduced pressure to give the crude intermediate **17**.

To a solution of **17** (145.2 mg, 1 mmol) in methanol (3 mL) was added SOCl_2_ (130.9 mg, 1.1 mmol) dropwise under ice bath. The solution was stirred at room temperature for 4h, and then the mixture was concentrated under reduced pressure. The residue was further purified by column chromatography to give intermediate **18** (94.3 mg, 76%).

To a solution of triphenylphosphine (112.0 mg, 0.416 mmol) in THF (10 mL) was added DIAD (84.1 mg, 0.416) under ice bath. The reaction was stirred at 0 °C for 0.5h under N_2_ atmosphere, and then a solution of **18** (94.3 mg, 0.32 mmol) in THF (4 mL) was added. The mixture was stirred at 0 °C for 0.5 h. After that, **S4** (211.4 mg, 0.64 mmol) was added. The reaction mixture was allowed to warm to room temperature by stirring overnight. The mixture was concentrated under reduced pressure, and the residue was purified by column chromatography to give intermediate **19** (167.8 mg, 91%).,

Intermediate **19** (167.8 mg, 0.29 mmol) was dissolved in THF (3.5 mL), MeOH (1.2 mL) and H_2_O (1.2 mL). 1N LiOH (1.72 mL) was added to the reaction mixture slowly at 0 °C. The mixture was stirred at room temperature overnight. The mixture was acidified with HCl (1N) to pH=2 at 0 °C. The acidified mixture was extracted with ethyl acetate (3 × 10 mL), and the combined organic phase was washed with brine, dried over anhydrous Na_2_SO_4_ and concentrated under reduced pressure to give intermediate **20** (152.4 mg, 93%).

To a solution of **20** (101 mg, 0.18 mmol) and **15** (106.9 mg, 0.27 mmol) in DMF (4 mL) was added HATU (136.9 mg, 0.36 mmol) and DIPEA (116.3 mg, 0.9 mmol). The reaction mixture was stirred at room temperature overnight. The mixture was concentrated under reduced pressure and was further purified by column chromatography to get intermediate **21** (110.2 mg, 72%).

To a solution of **20** (101 mg, 0.18 mmol) and **16** (119.0 mg, 0.27 mmol) in DMF (4 mL) was added HATU (136.9 mg, 0.36 mmol) and DIPEA (116.3 mg, 0.9 mmol). The reaction mixture was stirred at room temperature overnight. The mixture was concentrated under reduced pressure and was further purified by column chromatography to get intermediate **22** (100.7 mg, 63%).

To a solution of **21** (110.2 mg, 0.13 mmol) in methanol (2 mL) was added palladium hydroxide on activated carbon (10 mg, 10%). The mixture was stirred under H_2_ atmosphere at room temperature for 3h. The mixture was filtered, and the filtrate concentrated under reduced pressure. The residue was further purified by column chromatography to get intermediate **23** (84.2 mg, 92%).

To a solution of **22** (100.7 mg, 0.11 mmol) in methanol (2 mL) was added palladium hydroxide on activated carbon (10 mg, 10%). The mixture was stirred under H_2_ atmosphere at room temperature for 3h. The mixture was filtered, and the filtrate concentrated under reduced pressure. The residue was further purified by column chromatography to get intermediate **24** (97.9 mg, 100%).

To a solution of **23** (84.2 mg, 0.12 mmol) in THF (1.5 mL) was added H_2_O (2 drops) and DIPEA (41.9 mg, 0.32 mmol). Acryloyl chloride (10.9 mg, 0.12 mmol) was added dropwise in an ice bath. The mixture was allowed to warm to room temperature and stirred overnight. The solution was concentrated under reduced pressure, and the residue was further purified by column chromatography to give **IB-C5-DTB** (25.6 mg, 28%).

To a solution of **24** (97.9 mg, 0.13 mmol) in THF (1.5 mL) was added H_2_O (2 drops) and DIPEA (46.7 mg, 0.36 mmol). Acryloyl chloride (12.1 mg, 0.13 mmol) was added dropwise in an ice bath. The mixture was allowed to warm to room temperature and stirred overnight. The solution was concentrated under reduced pressure, and the residue was further purified by column chromatography to give **IB-C8-DTB** (51.1mg, 49%).

### Reagents & Cell culture

Spebrutinib and ibrutinib were purchased from MedChemExpress. Unless otherwise stated, all the chemicals were purchased from Sigma Aldrich. Ramos and Hela cell line was purchased from American Type Culture Collection (ATCC, Cat. No. CRL-1596 and CCL-2). Ramos cells were maintained in RPMI-1640 (Corning, Cat. No. 10-040-CV) supplemented with 10% fetal bovine serum (FBS) (Corning, Cat. No. 35-011-CV). Hela cells were cultured in DMEM (Corning, Cat. No. 10-013-CV) supplemented with 10% FBS. All the cell cultures were maintained at 37°C with 5% CO_2_. Cells were used for COOKIE experiments only when the cell viability is higher than 95%, confirmed by trypan blue exclusion assay. The 5x NP-40 cell lysis buffer contains 250 mM Tris-HCl pH 7.5, 750 mM NaCl, 2.5% NP-40 Alternative (Millipore, Cat. No. 492018), 50% glycerol, 5 mM EDTA. Halt™ Protease inhibitor cocktail 100x (Thermo Scientific, Cat. No. 78440) was freshly added into the 5x NP-40 lysis buffer to 5x final concentration before the COOKIE experiment.

### COOKIE-Pro protocol for advanced compound off-target kinetics profiling COOKIE sample preparation for immunoblotting confirmation

Ramos cells were cultured until they met the number requirement of COOKIE experiment. Each dose and timepoint sample require 2 million Ramos cells (equal to ∼120 ug total protein). In our COOKIE-Pro design, a total of 17 samples needs at least 34 million cells. To collect the cells, Ramos cell culture was centrifuged at 500 x g for 5 min followed by aspiration of the culture media. Cell pellet was washed and resuspended in phosphate buffered saline (PBS) pH 7.4 (Corning, Cat. No. 21-040-CV) to make the final cell density 10 million/mL.

In the first “pulse” step, for DMSO control sample, 2 million cells in 200 µL were incubated with 0.005% digitonin followed by 1 min 1% DMSO incubation. For saturating control sample, 2 million cells in 200 µL were incubated with 0.005% digitonin followed by 5 µM spebrutinib or ibrutinib incubation for 60 min. For each drug concentration, 6 million cells (for 3 timepoints) in 600 µL volume were incubated with 0.005% digitonin followed by adding 6 µL 100x concentrated spebrutinib or ibrutinib dissolved in DMSO for 2/4/6 min incubation. To start the second “chase” incubation, at the corresponding “pulse” timepoint, 180 µL permeabilized cell suspension was withdrawn and transferred into 45 µL 5x NP-40 lysis buffer containing 25 µM SB-2 or 5 µM IB-C8-DTB with a short gentle vortexing. The second “chase” incubation lasts for 1 hour for all samples. Both incubation steps were under room temperature.

After chase incubation, all the samples were desalted to remove free DTB probes using Zeba™ Spin Desalting Plates (Thermo Scientific, Cat. No. 89807). Although the sample volume has exceeded the recommended Zeba plate loading volume, we confirmed that this condition does not impair the pulldown yield (**Figure S6A, B**). For denature pulldown protocol, 0.1% SDS was added to the desalted samples. Each desalted sample was then incubated with 50 µL slurry Dynabeads™ M-280 Streptavidin (Invitrogen, Cat. No. 11206D) pre-equilibrated three times with 1x NP-40 lysis buffer with 0.1% SDS. After overnight bead binding at 4°C with rotation, the beads were separated from the lysate with magnetic rack. Lysate was aspirated and the beads were consecutively washed with 1x NP-40 lysis buffer twice and fresh-made 8 M urea twice. The beads were then transferred into new tubes and further washed with PBS twice. The beads were finally resuspended in 40 µL 200 mM HEPES pH 8.5. To prepare the immunoblot samples, a 5 µL aliquot was taken from each slurry bead sample and mixed with 1.6 µL 4x Laemmli buffer (Bio-Rad, Cat. No. 1610747) containing 10% β-mercaptoethanol. The immunoblot samples were incubated at 95°C for 5 min and ready to load onto SDS-PAGE gels. The remaining slurry bead samples were stored at -20°C until immunoblotting confirmed successful pulldown.

### TMT-labeled proteomics sample preparation

The 35 µL slurry bead samples were thawed, resuspended, and reduced with 5 mM dithiothreitol (DTT) (Fisher Scientific, BP172) at 37°C for 15 min with vigorous shaking, followed by alkylation with 20 mM iodoacetamide dissolved in 200 mM HEPES pH 8.5 (Thermo Scientific, Cat. No. A39271) at room temperature in the dark for 30 min. After aspirating all the supernatant, on-bead digestion was initiated by resuspending the beads in 40 µL 12.5 ng/uL trypsin/LysC mix (Promega, V5073) diluted in 200 mM HEPES pH 8.5. The bead-protease suspension was vigorously shaken (1,100 rpm) on a 37°C tube incubator for 18 hours. Acetonitrile (ACN) was then added to the digested samples to reach 30% v/v. For each sample, one TMTpro-18plex channel was assigned, and 10 µL of 25 µg/µL corresponding TMTpro reagent (Thermo Scientific, Cat. No. A52045), dissolved in 100% ACN, was added and incubated for 1 hour at room temperature. The labeling reaction was quenched by adding 10 μL of 5% hydroxylamine to each sample and incubating for 15 min. All the samples were combined, acidified with formic acid (FA) until pH < 3, and dried in the SpeedVac. The dried sample was reconstituted in 100 µL 200 mM HEPES pH 8.5 and desalted using SP3 peptide clean-up protocol ^10,49^. Briefly, 200 ug 1:1 mixed Sera-Mag Carboxylate SpeedBeads E3 and E7 (Cytiva # 65152105050250 and # 45152105050250) were prewashed and added into the peptide sample. The binding process was initiated by adding 1.8 mL 100% ACN with vigorous vortex followed by 10 min 1,000 rpm shaking. The supernatant was removed, and the beads were washed and vortexed twice with 200 µL 100% ACN. The beads were air-dried, and peptides were eluted using 2% DMSO for 10 min with intermittent sonication. Desalted peptides were SpeedVac dried until ready to inject into nanoLC-MS/MS.

### Data dependent acquisition (DDA) on nanoLC-MS/MS

The peptides were reconstituted in 5% MeOH/0.1% FA. Each sample was first separated by a 15 cm PepMap™ 100 C18 HPLC Column (Thermo Scientific, Cat. No. 164534) on Thermo EASY-nLC through a 5–30% B gradient for 95 min followed by a 30-60% B gradient for 38 min (B = 90% ACN) at 300 nL/min flow rate. The peptides were eluted into an Orbitrap Fusion Lumos mass spectrometer (Thermo Fisher Scientific). Peptides were ionized by electrospray (+2.4 kV), followed by MS/MS in data-dependent acquisition mode with a 3 sec cycle-time. MS1 data were acquired using the orbitrap analyzer in profile mode at a resolution of 120,000 over a range of 400–1,600 m/z at 100% normalized automatic gain control (AGC). Following HCD at 38% normalized collision energy and quadrupole isolation with a window of 1.2 m/z, MS2 data were acquired using the orbitrap at a resolution of 50,000 in centroid mode with the first mass 110 m/z at 200% normalized AGC.

### COOKIE-Pro data analysis

Proteome Discoverer 2.4 (Thermo Fisher Scientific) was used for RAW file processing, controlling peptide- and protein-level false discovery rates, assembling proteins from peptides, and protein quantification from peptides. Searches were performed using full tryptic digestion against the SwissProt human database with up to two missed cleavage sites. Oxidation (+15.995 Da) on methionine and deamidation on N and Q (+0.984 Da) were set as variable modifications, while carbamidomethylation of cysteine residues (+57.021 Da) and TMTpro labeling of peptide N-termini and lysine residues were set as fixed modifications (+304.207 Da). Data were searched with mass tolerances of ± 20 ppm and 0.5 Da on the precursor ions and fragment ions, respectively. The results were filtered to include peptide spectrum matches (PSMs) with a high peptide confidence with PSM and peptide-level FDR<0.01. PSMs with precursor isolation interference values > 50% and average TMT-reporter ion signal-to-noise values (S/N) < 10 were excluded from quantitation. Isotopic impurity correction and TMT channel normalization, based on the total peptide amount, were applied. Protein quantification uses both unique and razor peptides, with protein-level FDR <0.01 for strict and <0.05 for relaxed restrictions.

Reported protein abundance for each TMT channel was processed for COOKIE two-step fitting in Prism v10.2 (GraphPad Software). Each individual protein abundance was first normalized against acetyl-CoA carboxylase 1 (ACC1) among all the samples. Then for each individual protein, the ACC1-normalized abundance was converted to control sample-normalized abundance by setting 0% as the protein abundance in the saturating control sample, and 100% as the abundance in the DMSO control sample. This control sample-normalized abundance is equal to the occupancy of DTB probe on each individual protein, and the occupancy of testing inhibitor 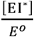 can be calculated from (1 - occupancy of the DTB probe). With 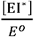 at different concentrations and timepoints, *k*_*obs*_ can be fitted for each concentration using the one-phase association model with *Y* = *Y*0 + (*Plateau Y*0) ∗ (1 − *exp* (− (*k*_*obs)* ∗ *x*)), where *x* = *time* in seconds, 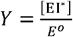, the *Plateau* is constrained to 1 and the *Y*0 constrained to 0. The ([*I*],*k*_*obs*_) pairs were further plotted for the second Michaelis-Menton fitting using user-defined model *Y*= *kinact* ∗ *X* /(*KI* + *X*), where the initial value of *kinact* set as 1*YMAX and *KI* set as 1*(Value of X at YMID).

### Two-point COOKIE-Pro protocol for covalent fragment screening

#### Two-point COOKIE-Pro sample preparation with SP3 protein clean-up, TMT labeling, and peptide-level streptavidin enrichment

A day before the experiment, 5 million Hela cells were plated in a 10 cm cell culture dish with ∼80% confluency. This amount of cell generates ∼1.5mg total protein for 4 replications of TMTpro 18plex experiments with 20 μg total protein per TMT channel. After overnight culture, cells were washed with PBS and lyzed in the PBS lysis buffer pH 7.4 containing 0.1% NP-40, 1x Halt™ Protease inhibitor cocktail (Thermo Scientific, Cat. No. 78440), and 5 μL Benzonase (Sigma-Aldrich, E1014) for 10 minutes. The lysate was centrifuged at 14,000 ×g for 5 min, and the supernatant was normalized to 2 μg/μL protein concentration. A total of 10 μL lysate (20 μg total protein) was dispensed into each well of a 96-well deep plate. Covalent fragments stocks in DMSO were diluted as 3x in the lysis buffer beforehand. A volume of 5 μL fragment were added into each well. For each fragment, two concentrations, 20 μM and 50 μM were used to incubate with the lysate for 1 hour under room temperature. Desthiobiotin-iodoacetamide (DBIA) probe (2 mM in lysis buffer, 5 μL) were added into each well with a final concentration of 500 μM to react for another 1 hour in dark.

After incubation, the reaction mix was cleaned-up using SP3 protein clean-up protocol ^49,50^. Briefly, 150 μg of SP3 beads (1:1 mixture of E3 and E7 carboxylic magnetic beads from Cytiva Cat. #45152105050250 and Cat. #65152105050250) were added into each well with a brief vortex to resuspend beads. A total of 30 μL of 20 mM DTT dissolved in 100% ethanol was added into each well, followed by 10 min 1,000 rpm shaking to trigger protein binding to the beads. Supernatant was aspirated and another 20ul aliquot of lysis buffer containing 20 mM iodoacetamide was added, and the plate was vigorously shaken for 30 min in dark. Supernatant was aspirated, and beads were washed twice with 60 μL 80% ethanol. The beads were finally resuspended in 30 μL 12.5 ng/μL trypsin/LysC (Promega, V5073) in 200 mM HEPES pH 8.5. The plate was filmed and incubated at 37°C overnight with 1,000 rpm shaking.

After digestion, 11 μL ACN were added into the product peptide-bead mixture. To each sample, 120 μg TMTpro 18-plex reagent freshly dissolved in 4 μL ACN was added and incubated at room temperature for 1 hour. The reaction was quenched by adding 7ul 5% hydroxylamine for 10 min. Samples with different TMTpro channel labeling were combined, with beads removed by magnetic stand, and SpeedVac to dry. Peptides were resuspended in H_2_O, acidified with formic acid, and further cleaned up with SepPak SPE C18 column using 5% ACN/0.1% FA (Waters, WAT054955). Eluted peptides were dried, resuspended in 300 μL HEPES pH 8.5 buffer, and DBIA-labeled peptides were enriched using 50 μL High Capacity Streptavidin Agarose (Pierce, #20357) with 3 h end-to-end rotation at room temperature. Agarose beads were washed in the filter unit with twice HEPES/0.05% NP-40 pH 8.5 buffer, three times HEPES pH 8.5 buffer, and once H_2_O. Bound DBIA-peptides were eluted with 80% ACN/0.1% FA/0.2% TFA with three consecutive incubation with shaking 1) RT 20min, 2) RT 10min, 3) 72°C 10min ^10^. Eluted fractions were combined, speedvac dried, and further cleaned up with C18 StageTips. Dried peptides were kept in -20°C before LC-MS/MS analysis.

#### Data dependent acquisition (DDA) of DBIA peptides on nanoLC-FAIMS-MS/MS

Dried peptides were resuspended in 5% ACN/0.1% FA and each sample injected twice as technical replicates. Peptide samples were separated and analyzed using a Vanquish Neo UHPLC system coupled to an Orbitrap Ascend mass spectrometer (Thermo Fisher Scientific). The UHPLC was configured for direct injection with a 50 cm * 75 µm inner diameter separation column. Mobile phase A consisted of H_2_O/0.1% FA and mobile phase B consisted of 80% ACN/0.1% FA. Peptides were separated using a 150-minute gradient. The flow rate was maintained at 0.250 µL/min while the percentage of mobile phase B was increased from 4% to 25% over 120 minutes, then to 40% B from 120 to 140 minutes. The column was subsequently washed by increasing the flow rate to 0.350 µL/min and the concentration of mobile phase B to 99% from 141 to 150 minutes.

The Orbitrap Ascend mass spectrometer was operated in positive ion mode and utilized High-Field Asymmetric Waveform Ion Mobility Spectrometry (FAIMS) with standard resolution. The analysis was conducted by cycling through two FAIMS compensation voltages (CVs): -50 V and -70 V. The total cycle time for the data-dependent acquisition was 1.5 seconds.

Full MS1 scans were acquired in the Orbitrap at a resolution of 120,000 over a scan range of m/z 400-1600 (for CV -50) and m/z 400-1500 (for CV -70). The maximum injection time for MS1 scans was set to 25 ms for the -50 CV and Auto for the -70 CV. The normalized Automatic Gain Control (AGC) target was 50% (absolute target of 2e5) for the -50 CV experiment and 25% (absolute target of 1e5) for the -70 CV experiment.

For MS2 analysis, precursor ions with charge states from 2 to 6 and a minimum intensity of 5e3 were selected for fragmentation. Dynamic exclusion was enabled to exclude precursors after one acquisition for a duration of 45 seconds with a mass tolerance of 10 ppm. Selected precursors were isolated using the quadrupole with an isolation window of 1.0 m/z and fragmented via Higher-energy Collisional Dissociation (HCD) with a normalized collision energy of 38%. The resulting fragment ions were detected in the Orbitrap at a resolution of 30,000. The MS2 scans had a first mass fixed at m/z 120. The maximum injection time for MS2 scans was set to 96 ms, with a normalized AGC target of 200% (absolute target of 1e5). Data was acquired in centroid mode for MS2 scans.

#### Two-point COOKIE-Pro data analysis

The raw data files were processed using Proteome Discoverer™ software (version 3.2, Thermo Fisher Scientific). All MS/MS spectra were searched against the UniProt *Homo sapiens* reviewed proteome database (UP000005640, downloaded on January 7, 2025) using the Sequest HT search algorithm. A target-decoy strategy was employed to filter peptide-spectrum matches (PSMs) to a 1% false discovery rate (FDR) using the Percolator node.The search parameters for the fully tryptic digestion allowed for a maximum of two missed cleavages. A precursor mass tolerance of 20 ppm and a fragment ion mass tolerance of 0.6 Da were applied. Tandem mass tags (TMTpro) labeling on peptide N-termini and lysine (K) residues (+304.207 Da) were set as static modifications. The following dynamic modifications were permitted: Oxidation of methionine (M, +15.995 Da), Deamidation of asparagine (N) and glutamine (Q) (+0.984 Da), Carbamidomethyl (+57.021 Da) and DBIA (+296.185 Da) on cysteine (C). Quantification of TMTpro reporter ions was performed using the Reporter Ions Quantifier node based on the HCD MS2 scans. The reporter ion intensities were quantified using the “Most Confident Centroid” method with an integration tolerance of 20 ppm. PSMs with precursor isolation interference values > 50% and average TMT-reporter ion signal-to-noise values (S/N) < 10 were excluded from quantitation. Isotopic impurity correction and TMT channel normalization, based on the total peptide amount, were applied. Protein quantification uses both unique and razor peptides.

Peptide group search results were exported, and further COOKIE calculation was processed with a custom

Python script. For the high-throughput covalent fragment screening performed at a single time point with two concentrations, a specialized analysis was employed to classify the kinetic behavior of each fragment-peptide interaction. After calculating the *k*_*0bs*_ value for each of the two concentrations, three distinct kinetic models were fitted to the *k*_*0bs*_ data points:

Linear Model: *k*_*0bs*_ = (*k*_*inact*_/*K*_*I*_) _·_ [*I*], applicable when inhibitor concentrations [I] are much lower than KI. Hyperbolic Model: *k*_*0bs*_ = *k*_*inact* ·_ [*I*] /(*KI* + [*I*]), the full Michaelis-Menten model.

Constant Model: *k*_*0bs*_ = *k*_*inact*_, applicable when inhibitor concentrations [I] are much higher than *KI*.

To select the most appropriate model for each pair while avoiding overfitting, the Akaike Information Criterion (AIC) and Bayesian Information Criterion (BIC) were calculated for each successful model fit under the assumption of Gaussian independent and identically distributed residuals.

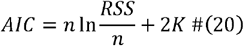

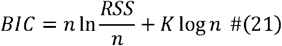

where 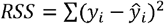, and *K* indicates the number of parameters to fit in each model. K= 1 in linear and constant model, K= 2 in hyperbolic model.

The script selects the model with the lowest AIC score as the best representation of the kinetic behavior.

This model selection process allows for the most reliable estimation of kinetic parameters from the limited two-point data, providing either the inactivation efficiency (*k*_*inact*_/*K*_*I*_) from the linear model, individual *k*_*inact*_ and *K*_*I*_ values from the hyperbolic model, or an estimate of *k*_*inact*_ and upper boundary of *K*_*I*_ from the constant model.

### Immunoblotting

The input sample before streptavidin pulldown and the boiled beads in the Laemmli buffer were separated by SDS-PAGE using 4-20% Criterion TGX precast gels (Bio-Rad, Cat. No. 5671095) and further semi-dry transferred onto 0.45 µm PVDF membrane (Millipore, Cat. No. IPVH85R) pre-soaked with ethanol. After 1 hour blocking with 4% bovine serum albumin in 1x TBST, the membranes were probed overnight at 4 °C with the specified primary antibodies at the dilution of 1:1000 (Cell Signaling Technology: BTK Rabbit mAb#8547; Abclonal: Tec Rabbit mAb A22074) or 1:2000 (β-actin Rabbit mAb#4970), followed by 3 times 10 min TBST washes and incubation with the HRP-conjugated secondary antibody (Kindle Biosciences, #R1006, 1:1000) for 1 hour at room temperature. For streptavidin blotting, after blocking, the membranes were probed for 1 hour with HRP-Conjugated Streptavidin (Thermo Scientific, Cat. No. N100) at the dilution of 1:10,000. After 3 times 10 min TBST washes, chemiluminescence was developed using SuperFemto ECL chemiluminescence kit (Vazyme, Cat. No. E423-01) and captured in the digital imaging system. Densitometry analysis was through KwikQuant Image Analyzer 5.9 software (Kindle Biosciences).

### BTK kinase activity assay

PhosphoSens-Kinetic Kinase Assays (AssayQuant Technologies, AQT0101) was used to monitor the inhibition kinetics of BTK by ibrunitib, IB-ac-DTB, IB-C5-DTB, IB-C8-DTB. Each well on the 384-well plate (Corning, 3701) contains 20ul mixture of 2.5 µL reaction buffer (500 mM HEPES pH 7.5, 0.1% Brij-35, 100 mM MgCl_2_), 2.5 µL 100 µM PhosphoSens Sensor peptide substrate, 2.5 µL 10 mM adenosine 5′-triphosphate disodium salt hydrate (ATP) solution, 2.5 µL 10 mM tris(2-carboxyethyl)phosphine hydrochloride (TCEP) solution, and 10 µL ddH_2_O. DMSO or the testing compounds dissolved in DMSO were directly dispensed into the well using Multidrop Pico 8 Digital dispenser (Thermo Scientific) giving a final concentration of 8.7 nM to 500 nM. With all the reagents mixed, 5 µL 6.4 nM His-BTK (SignalChem, B10-10H) in enzyme dilution buffer (20 mM HEPES pH 7.5, 0.01% Brij-35, 5% glycerol, and 1 mg/ml BSA) was added to achieve final 1.28 nM BTK. The plate was immediately filmed and fluorescence intensity (Ex. 360 nm Em. 492 nm) was recorded every 3 min for 3 hours on a SYNERGY H1 microplate reader (BioTek).

### GSH reactivity measured by LC-UV/MS

Test compounds at 1 mM were reacted with excess of glutathione (100 mM) in 100 mM HEPES / 20% ACN pH 7.5 at 37°C in singlicate. The compound reactivity was qualitatively measured through monitoring the loss of parent compound by integrating the extracted ion chromatogram and UV chromatogram on an Agilent 1260 Infinity II HPLC system tandemed with single quadrupole InfinityLab LC/MSD system.

### Zeba plate desalting capacity test

Zeba desalting plate volume capacity was initially tested with a desthiobiotin-BODIPY dye DTB-C6-BDP. Zeba plate was centrifuged at 1,000 x g for 2 min to remove the storage buffer and loaded with 120-320 µL 10 µM DTB-C6-BDP in PBS and centrifuged again to collect the desalted samples. The fluorescence intensities of the desalted samples were converted into concentrations through standard curve calibration. The standard COOKIE protocol was then tested with different sample volumes and cell numbers as in Figure S6B.

### Reprocessing SLC-ABPP dataset

We extracted the SLC-ABPP electrophile-peptide competition ratio (CR) values in HCT-116 cell screening from Kuljanin *et al*.^7^ Supplementary Table 6. In order to convert single concentration CR values into *K*_*I*_ values, we assumed that the *k*_*inact*_ reactivity should be similar for covalent warheads in the same class. The reported *k*_*inact*_ information was extracted from the CovalentInDB ^45^ and the unit was unified. After stratifying the entries into warhead subclasses, the *k*_*inact*_ median of chloroacetamide and acrylamide compounds were used for downstream calculation. The CR values greater than 2 were converted to *K*_*I*_ value through these three equations:

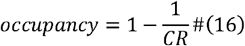

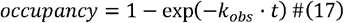

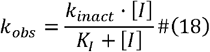

among which the *t*, and [*I*] was 7200 seconds and 2.5e-5 M from the SLC-ABPP method, *k*_inact_ for chloroacetamide was 0.0092 s^-1^ and for acrylamide was 0.0025 s^-1^. The ligand efficiency (LE) was calculated according to

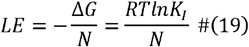

where Δ*G* is the molar Gibbs free energy, N is the atom number in the molecule. The ligand lipophilic efficiency (LLE) was calculated from:

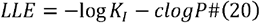

where the *clogp* was calculated using Crippen’s approach implemented in RDKit 2023.09 package. All the data analysis and visualization were achieved through Python 3.9.

### Statistics

For spebrutinib and ibrutinib COOKIE-Pro experiments, the *k*_*0bs*_ obtained from each independent COOKIE-Pro experiment were combined as replicates to perform the Michaelis-Menton fitting. The reported standard error of *k*_*inact*_ and *K*_*I*_ was calculated by GraphPad 10.2 using symmetrical approximate confidence intervals.

### Data reproducibility and code availability

COOKIE-Pro result for spebrutinib was reproduced in all three independent experiments. COOKIE-Pro result for ibrutinib was reproduced in all two independent experiments. Two-point COOKIE-Pro fragment screening was performed in two biological replicates, and each sample injected twice for LC-MS/MS analysis.

The mass spectrometry raw files for COOKIE-Pro quantitative multiplexed proteomics have been deposited in the MassIVE dataset under accession number MSV000095804. Python scripts for two-point COOKIE-Pro data analysis can be accessed through https://github.com/Hanfeng-Lin/COOKIE_scripts.

## Acknowledgments

This research was supported in part by the National Institutes of Health (R01-CA250503 and R01-CA268518 to J.W.), the Cancer Prevention & Research Institute of Texas (CPRIT, RP220480 to J.W.), and Michael E. DeBakey, M.D., Professor in Pharmacology (to J.W.).

## Supplementary Material

**Figure S1.**
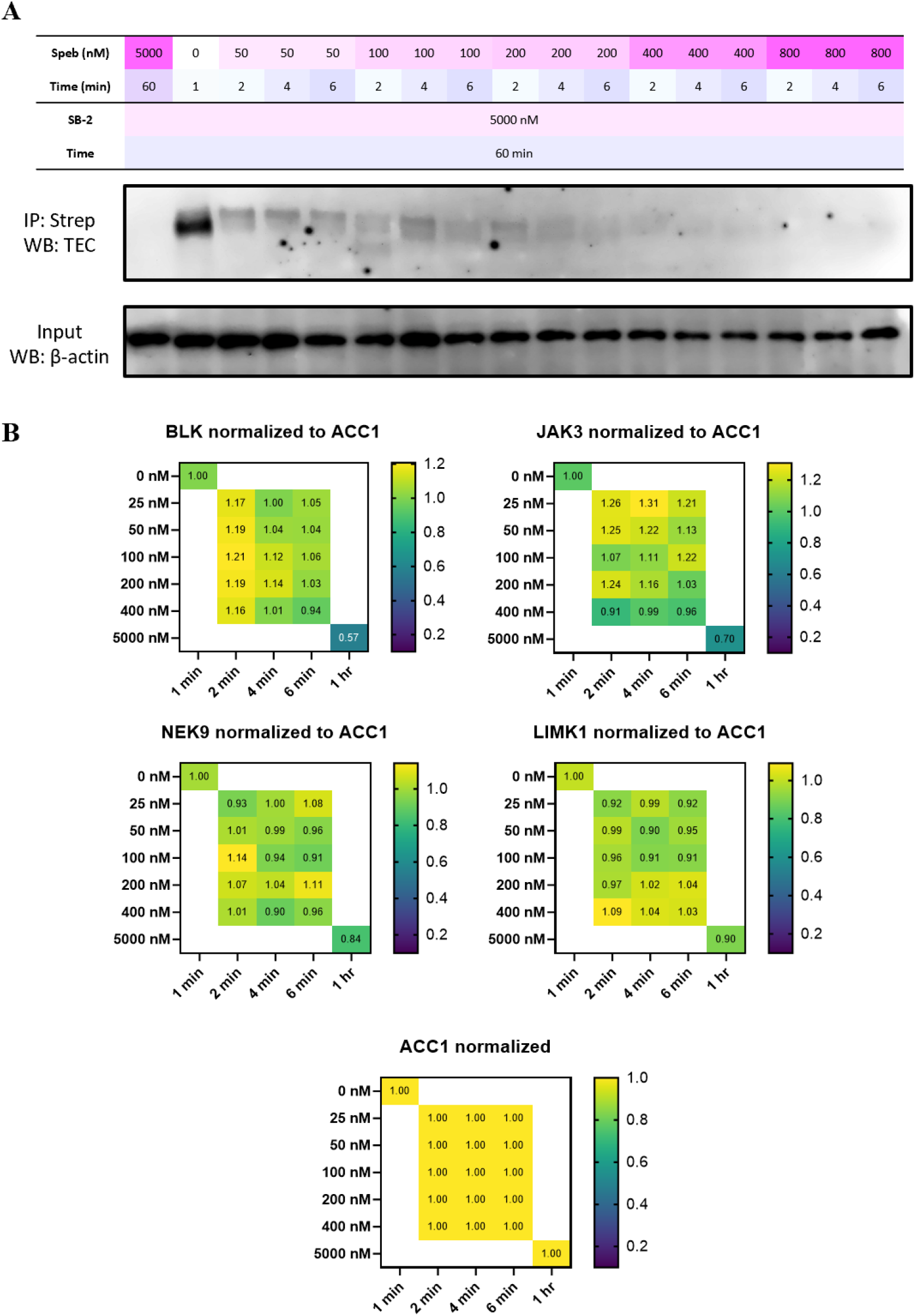
**(A)** Immunoblot of spebrutinib COOKIE samples indicating spebrutinib is a potent TEC kinase binder. **(B)** Spebrutinib off-targets that show engagement only at 5 µM treatment for 1 hour. All the data points have been normalized to the acetyl-CoA carboxylase 1 (ACC1) abundance.

**Figure S2.**
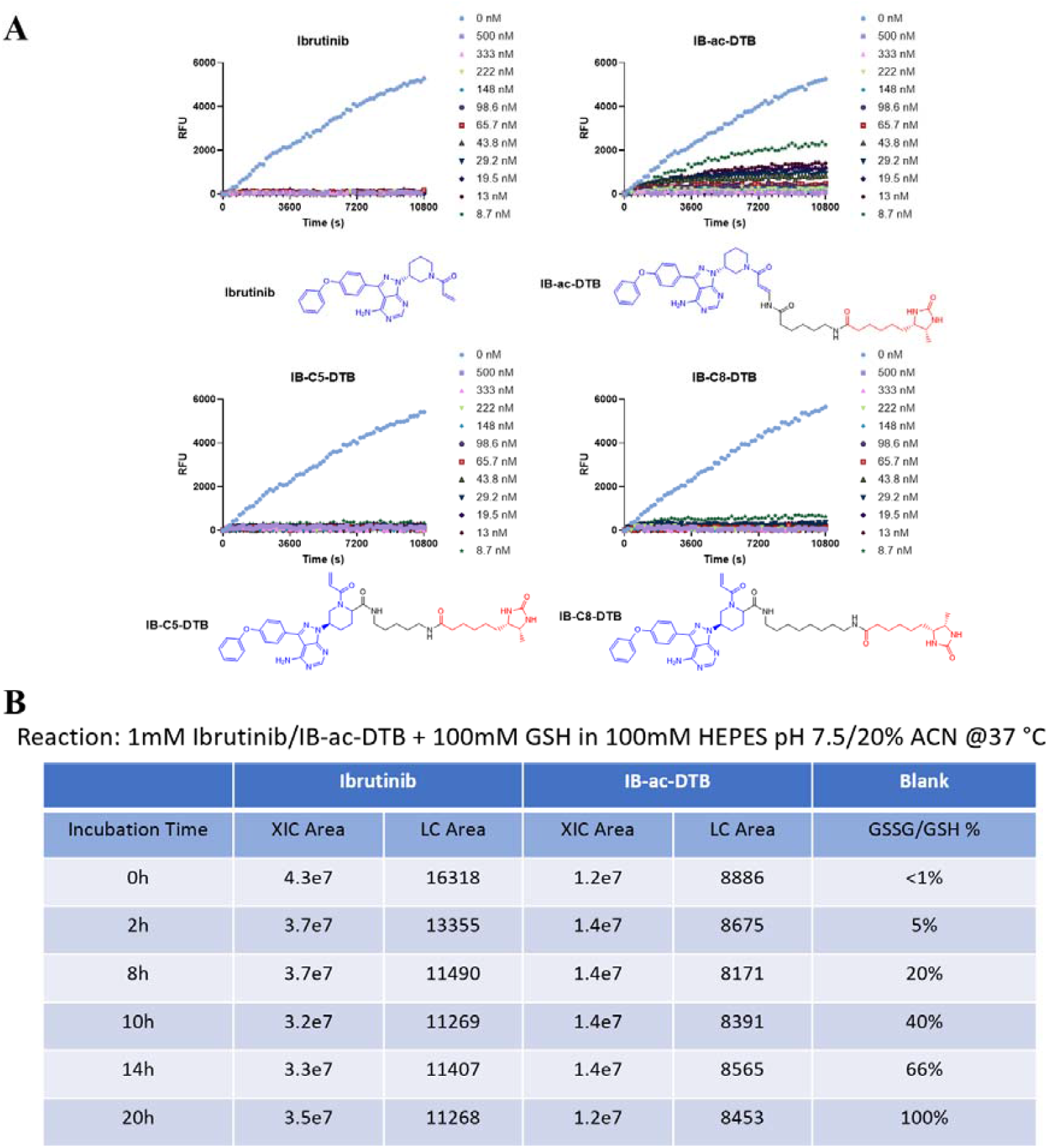
IB-DTB probes with different BTK inhibition potency and GSH reactivity using different exit vectors. **(A)** BTK kinase activity assay using PhosphoSens substrate and 1.28 nM BTK full-length protein. **(B)** GSH reactivity measured by LC-MS at intervals. IB-ac-DTB showed very weak GSH reactivity compared to ibrutinib. XIC, Extracted Ion Chromatogram.

**Figure S3.**
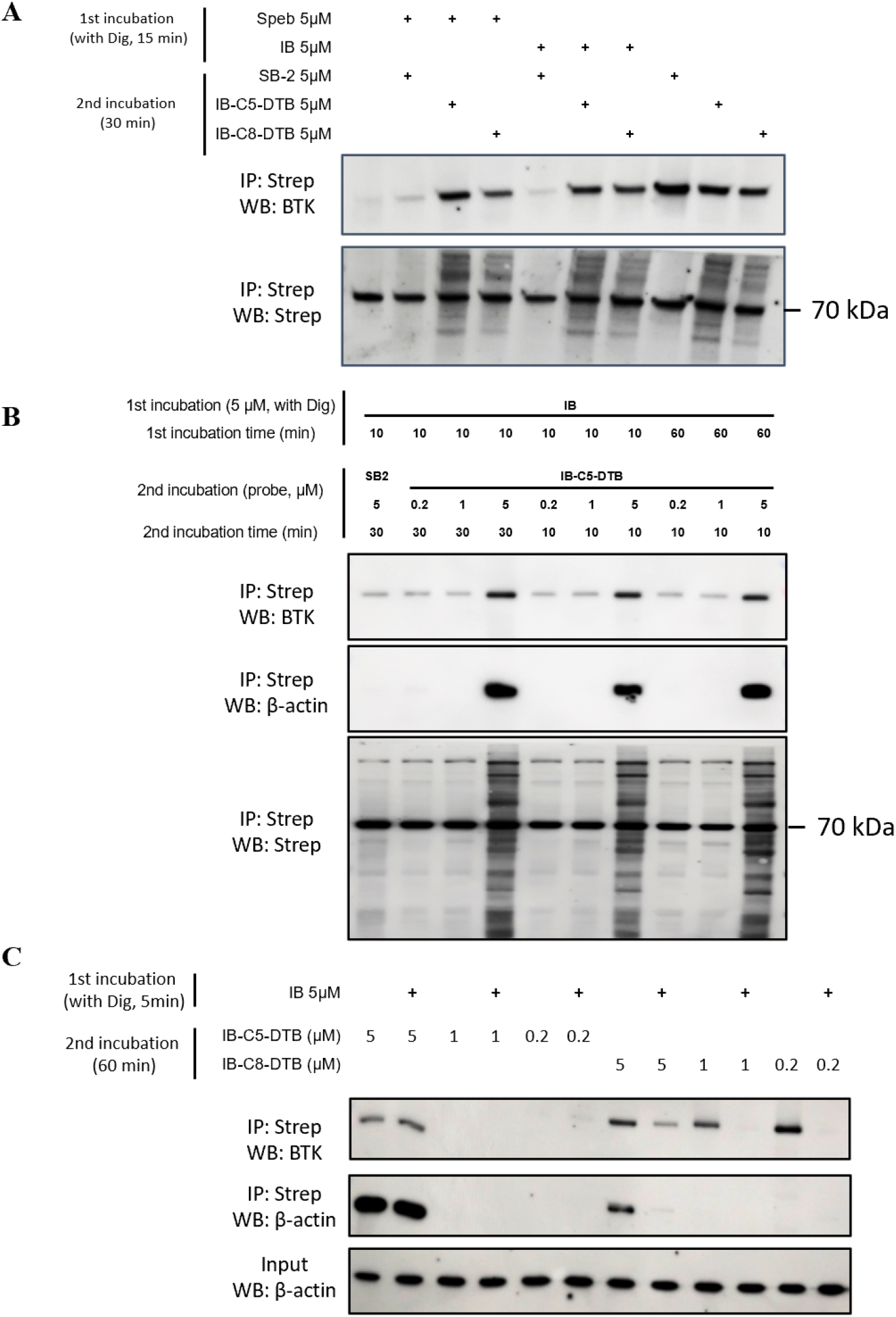
Optimizing the concentration of IB-DTB probe to avoid non-specific labeling and efficient pulldown. **(A)** IB-C5-DTB and IB-C8-DTB labeling at 5 µM cannot be competed out by either spebrutinib or ibrutinib. **(B)** IBC5DTB as low as 1 µM allows effective IB competition. **(C)** IB-C8-DTB, but not IB-C5-DTB, allows efficient pulldown under the native pulldown condition. IB, ibrutinib, Dig, digitonin.

**Figure S4.**
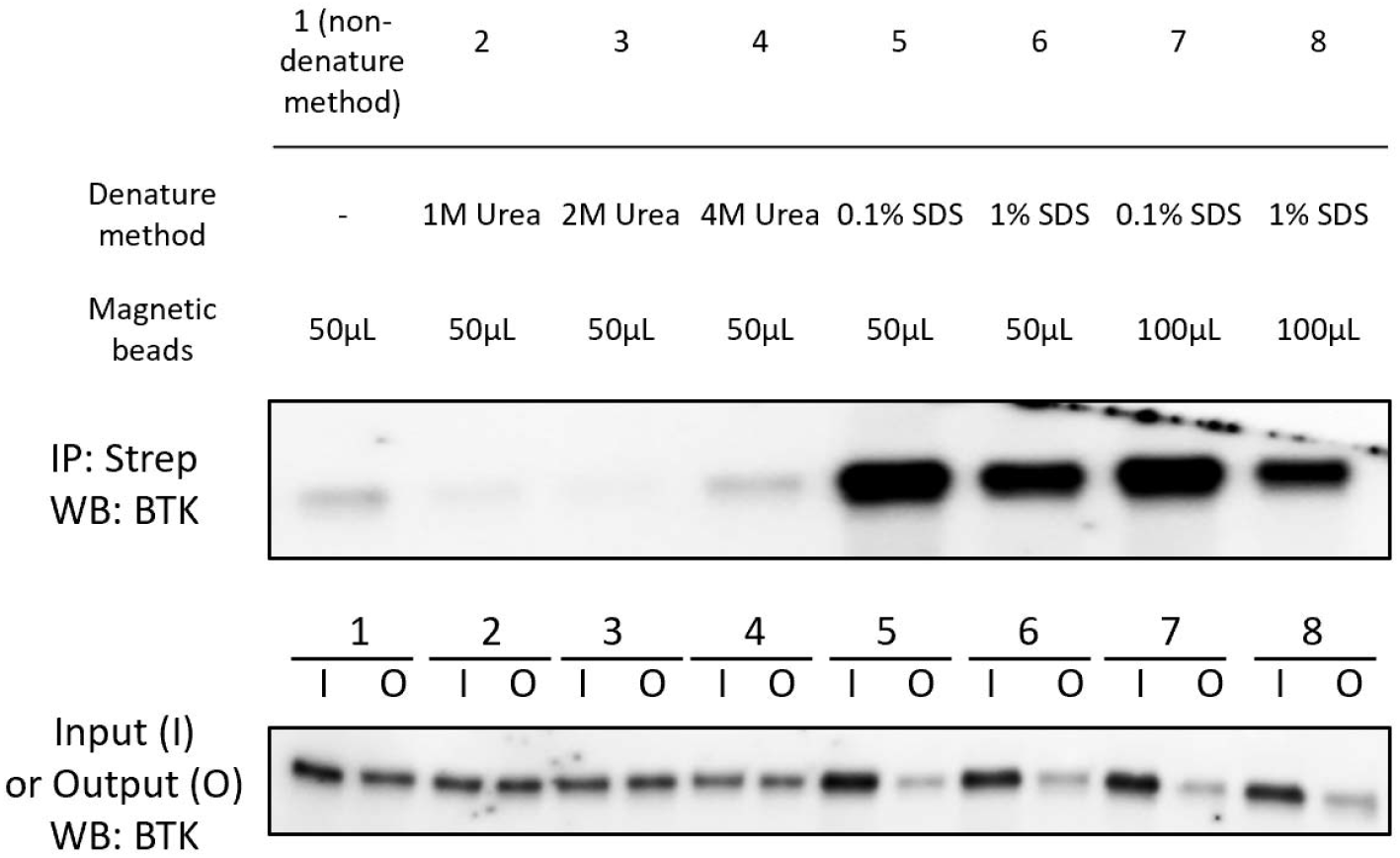
Denature pulldown condition optimization for IB-C8-DTB. A total of 2 million Ramos cells were used for each sample. Cells were lysed and incubated with 1 µ IB-C8-DTB for 1 hour at room temperature, followed by desalting and incubating overnight with indicated amount of streptavidin magnetic beads in the denature buffer containing urea or SDS.

**Figure S5.**
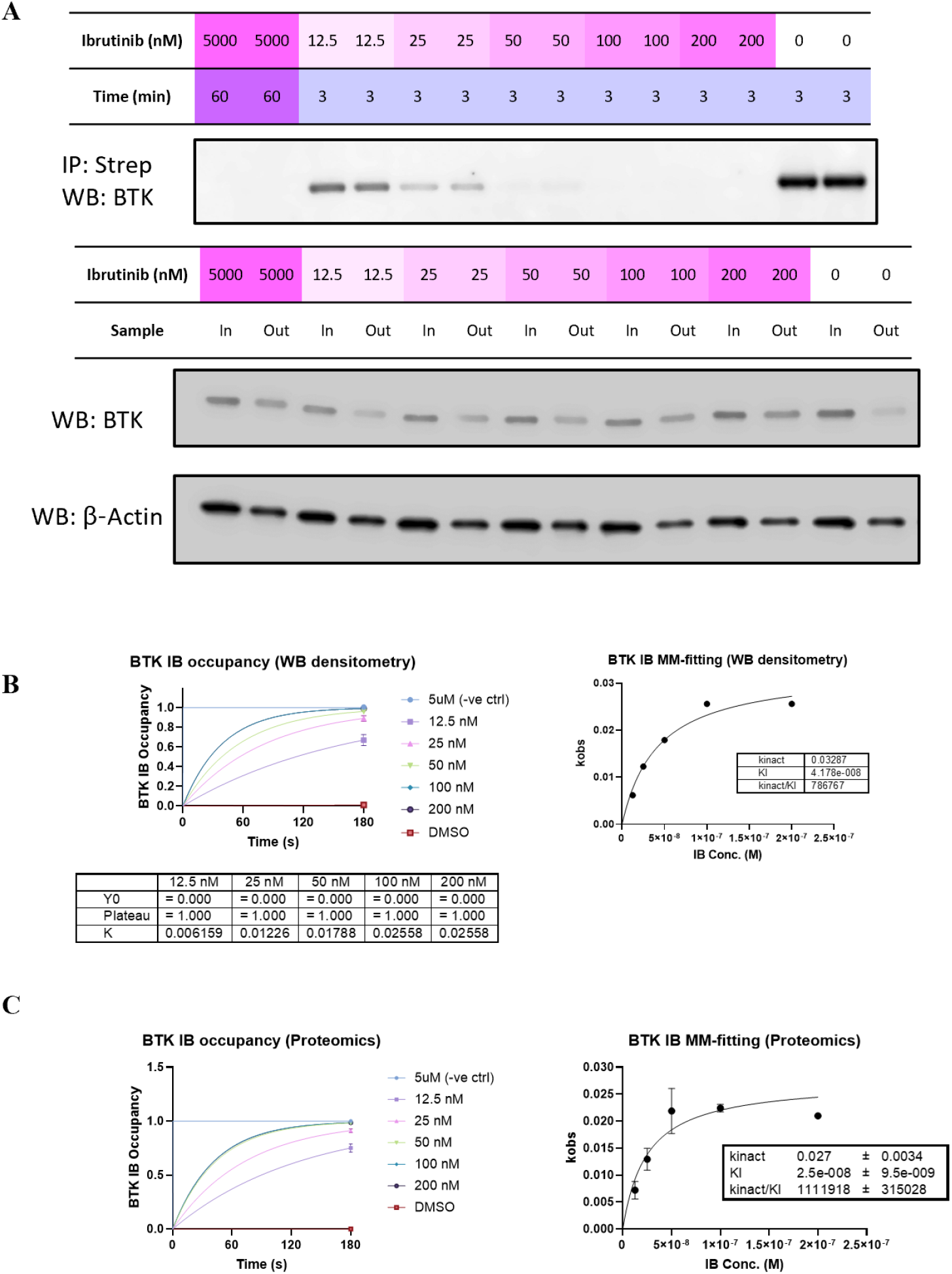
Single timepoint COOKIE profiling for ibrutinib in Ramos cell using IB-C8-DTB desthiobiotin probe. **(A)** Single timepoint ibrutinib COOKIE experiment layout and the immunoblots for BTK in the pulldown samples, input samples before pulldown and output samples after pulldown. **(B)** Densitometry analysis and (C) Proteomics analysis of panel A BTK pulldown sample blot plotted as first step progressive curve (left) and second step plot fitting into Michaelis-Menton model to obtain and values for ibrutinib against BTK. Data are presented as mean ± SD from two biological replicates.

**Figure S6.**
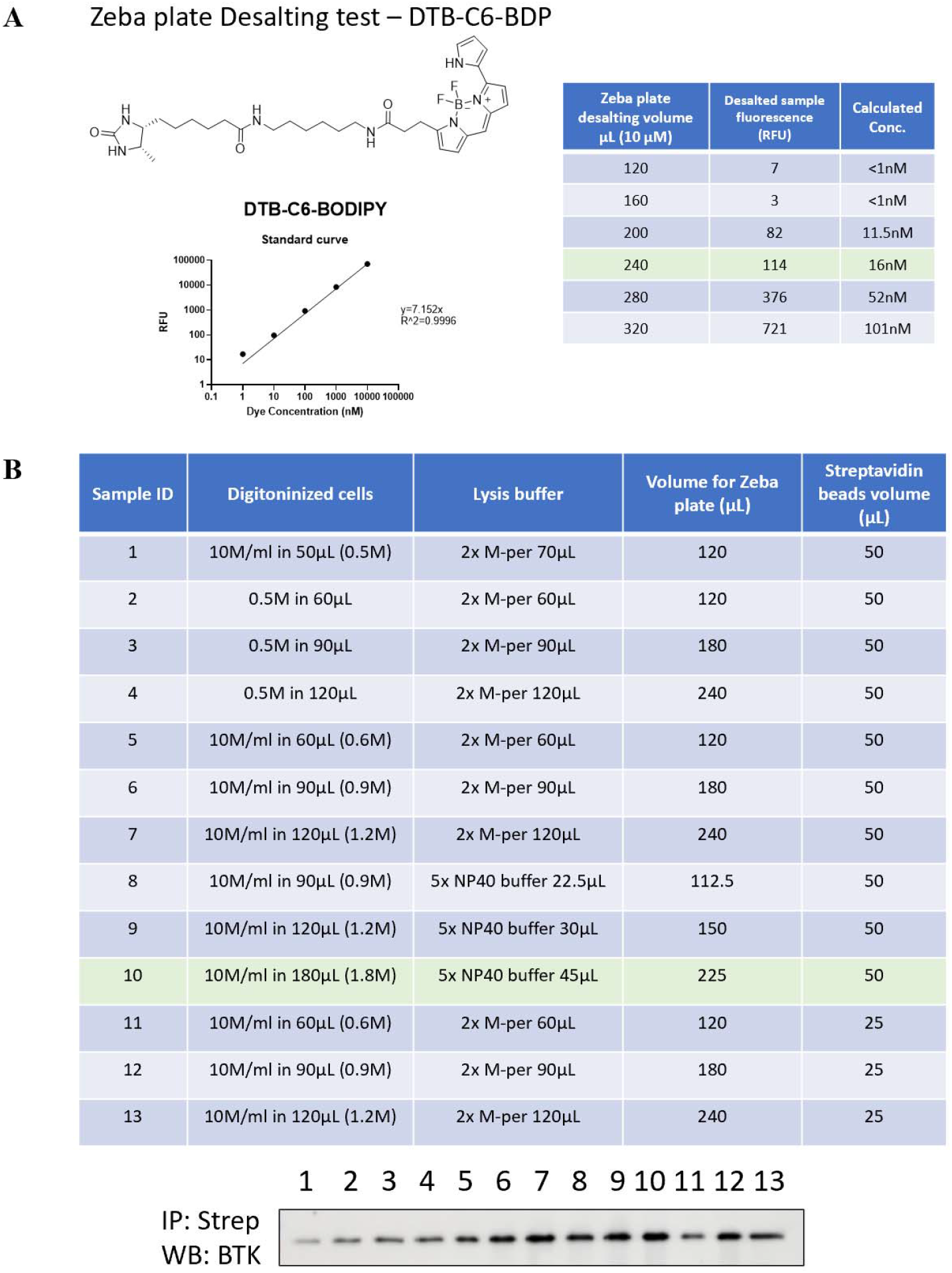
Desalting condition optimization to remove DTB probes. **(A)**. Off-label pressure test of Thermo Zeba desalting plate using increasing volume of 10 μM DTB-C6-BDP. A volume less than 200 μL can ensure 99.9% removal of small molecule dye in the sample. **(B)**. Multiple factor optimization for the pulldown protein amount reflected by the BTK WB band intensity from the IP sample.

**Figure S7.**
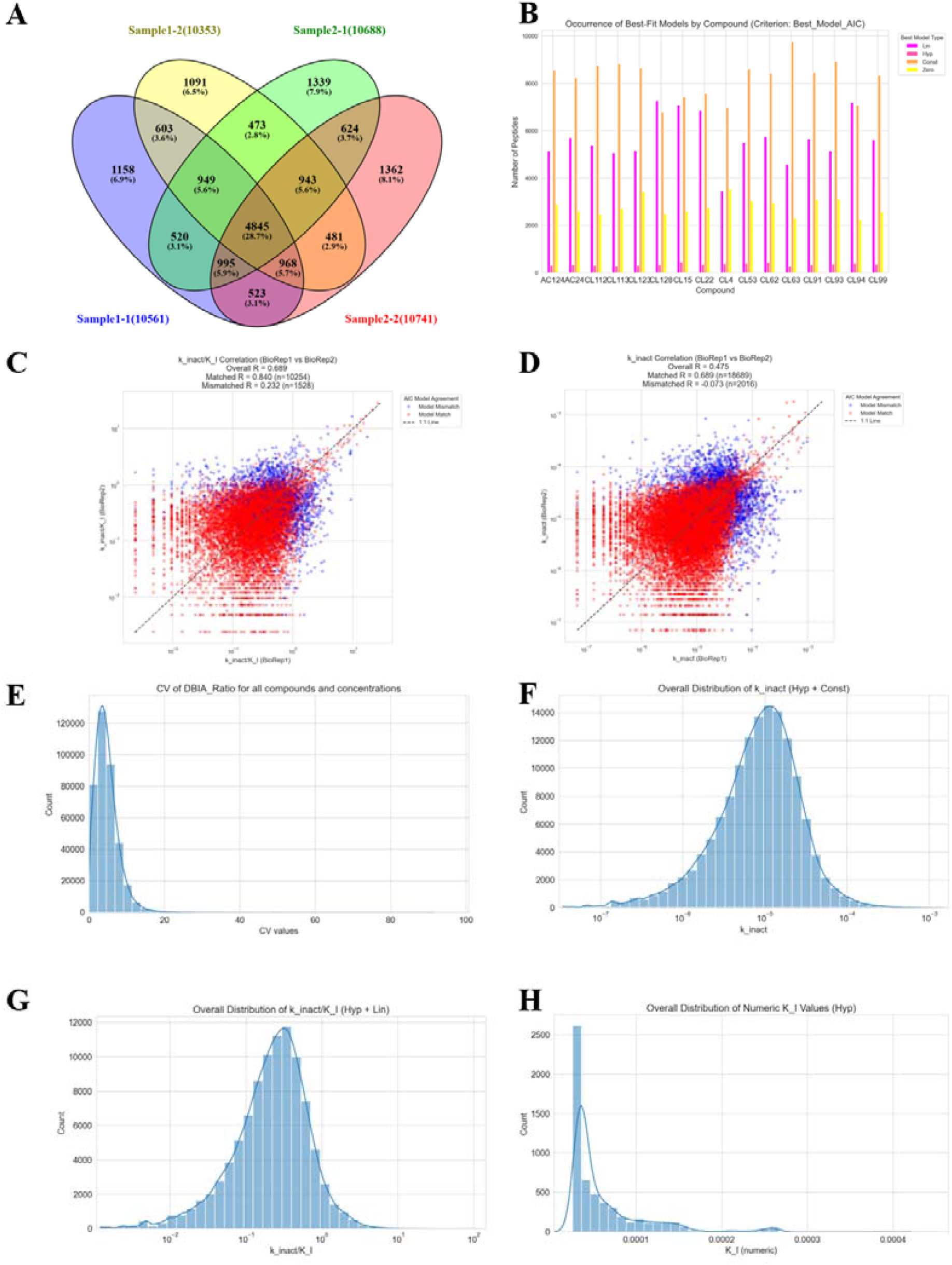
Descriptive statistics for two-point COOKIE-Pro fragment screening. **(A)** Venn diagram illustrating the overlap of identified DBIA-labeled peptide groups across two biological replicates (sample1, sample2) and two technical replicates each (e.g., sample1-1, sample1-2). **(B)** Bar chart showing the distribution of best-fit models (Zero, Linear, Hyperbolic, Constant) for each screened fragment. **(C)** Correlation plot of calculated inactivation efficiencies (*k*_*inact*_/*K*_*I*_) between two biological replicates, colored by matched/unmatched model selection for replicates. **(D)** Correlation plot of calculated reactivity (***k***_*inact*_) between two technical replicates, colored by matched/unmatched model selection for replicates. **(E)** Histogram showing the distribution of the coefficient of variation (CV) for all quantified DBIA peptide ratios across all conditions. **(F)** Histogram of the overall distribution of calculated ***k***_inact_) values derived from peptides fitting the hyperbolic and constant models. **(G)** Histogram of the overall distribution of calculated inactivation efficiencies (*k*_*inact*_/*K*_*I*_) from peptides fitting the hyperbolic and linear models. **(H)** Histogram of the overall distribution of calculated *K*_*I*_ values from peptides fitting the hyperbolic model.

